# Plant immune system activation is necessary for efficient interaction with auxin secreting beneficial bacteria

**DOI:** 10.1101/2021.03.23.436641

**Authors:** Elhanan Tzipilevich, Philip N. Benfey

**Affiliations:** Howard Hughes Medical Institute, Duke University, Durham, NC 27708, USA; Department of Biology, Duke University, Durham, NC 27708, USA

## Abstract

Plants continuously monitor the presence of microorganisms through their immune system to establish an adaptive response. Unlike immune recognition of pathogenic bacteria, how beneficial bacteria interact with the plant immune system is not well understood. Analysis of colonization of *Arabidopsis thaliana* by auxin producing beneficial bacteria revealed that activation of the plant immune system is necessary for efficient bacterial colonization and auxin secretion. A feedback loop is established in which bacterial colonization triggers an immune reaction and production of reactive oxygen species, which, in turn, stimulate auxin production by the bacteria. Auxin promotes bacterial survival and efficient root colonization, allowing the bacteria to inhibit fungal infection and promote plant health.

## Introduction

Plant roots interact with a plethora of bacteria in the surrounding soil. Extensive efforts have characterized the diversity of these bacterial species ^1, 2^. Bacteria can have pathogenic, beneficial or neutral effects on plants. Bacterial diversity is reduced when moving from bulk soil to the root surface (rhizosphere) and further into the root interior (endosphere), indicating that plants exert selective forces on their colonizing bacteria. The first filter utilized by plants to recognize and respond to bacteria and other organisms is its immune system ^3^, which utilizes receptors to recognize bacterial molecules called MAMPs (Microbe-Associated Molecular Patterns). These include flagella, peptidoglycans, bacterial elongation factor TU, and others ^4, 5^. Recognizing these molecules leads to a cascade of molecular events. At early stages these include an efflux of calcium ions and a burst of reactive oxygen species (ROS) ^6^. This is followed by phosphorylation events that lead to induction of immune-related genes ^7^. Plant immune system recognition and activation have been extensively characterized in the context of pathogenic bacteria ^8, 9^. However, MAMP receptors recognize molecules found in all bacteria, and beneficial bacteria can induce an immune response similar to pathogenic ones [e.g ^10^]. The influence of the immune system on the healthy root microbiome and how beneficial bacteria respond to the plant immune system is an active area of research and much remains to be learned ^11–13^. To better understand this process, we studied the interaction of *Bacillus valezensis FZB42* (*B. valezensis*) with the root of the model plant *Arabidopsis thaliana* (*Arabidopsis*). *B. valezensis* is a model gram-positive soil bacterium, which synthesizes a plethora of secondary metabolites shown to inhibit the growth of plant pathogens ^14^. It also synthesizes auxin ^15^, a plant hormone that influences many aspects of plant growth ^16^. A well characterized response of exogenous auxin addition is the arrest of primary root growth and stimulation of lateral root formation ^17^. *B. valezensis* was shown to stimulate lateral root formation and biomass accumulation in several plant species including *Arabidopsis*, *Lemna minor*, and lettuce, in an auxin-dependent manner ^18–20^. The effects of auxin secreted by bacteria on plant growth have been explored for decades. Bacterial auxin can manipulate plant growth ^21, 22^, probably providing the bacteria with access to nutrients. Bacterial auxin can also inhibit the plant immune system through antagonistic interaction with the salicylic acid signaling pathway ^23^. However, it is unclear if the production of auxin by bacteria has a direct effect on their colonization capacity ^24, 25^. We found that a positive feedback between the plant immune system and bacterial auxin secretion occurs during root colonization. Immune recognition of the bacteria triggers ROS production by the plant which in turn activates auxin secretion by the bacteria. This secreted auxin is necessary for bacterial survival in media containing elevated ROS levels, and for colony formation on the root. Efficient colony formation enables the bacteria to fight pathogenic fungi and enhance plant health. Thus, our work reveals that bacterial auxin directly impacts its capacity for root colonization, and uncovers a positive influence of the plant immune system on bacterial colonization with beneficial effects for the plant.

## Results

### Bacterial auxin plays a dual role during root colonization

To characterize the interaction between bacteria and plant roots, we inoculated *B. valzensis* onto Arabidopsis seedlings growing on agar plates. Consistent with previous results ^18, 20^, after 7 days of incubation, colonized plants exhibited reduced primary root growth and increased lateral root emergence (Fig 1A and Fig S1A-S1B) in comparison to seedlings treated with buffer (mock), a response consistent with bacterial auxin secretion [e.g ^22^]. Plants expressing an auxin response reporter [DR5::GFP ^26^] revealed increased auxin response in roots inoculated with *B. valzensis* (Fig 1B-1C). The root phenotype was dependent on bacterial auxin secretion, as plants inoculated with a strain of *B. valzensis* deficient in auxin production (*ΔysnE* ^18^) failed to exhibit primary root growth inhibition or lateral root stimulation (Fig 1A and Fig S1A-S1B) and had reduced DR5::GFP fluorescence (Fig 1B-1C).

**Fig 1.**
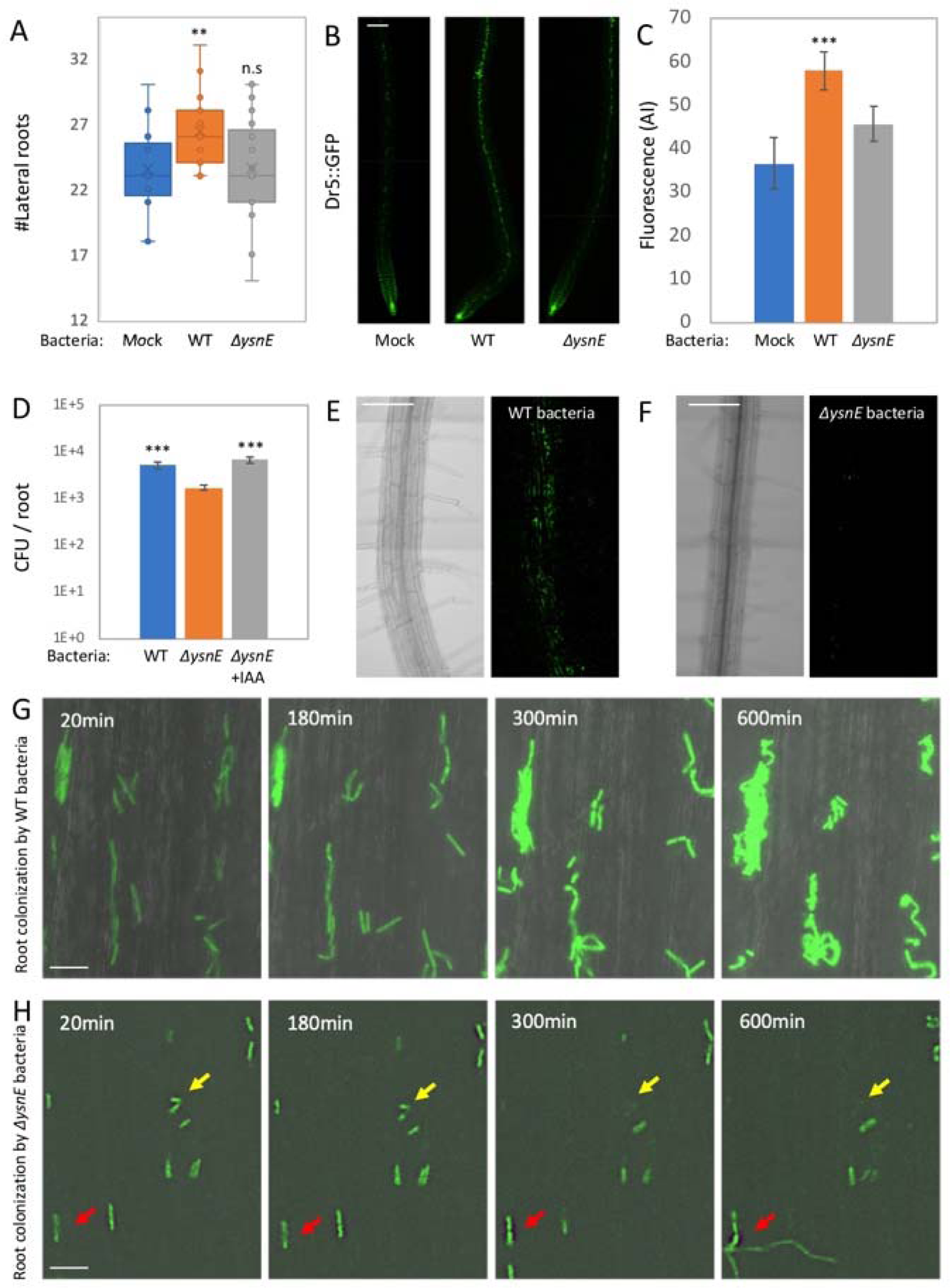
Bacterial auxin stimulates plant lateral root formation and bacterial root colonization. (A) Seedlings were inoculated with either WT, *ΔysnE* bacteria or buffer (mock) on agar plates for 7 days and the number of lateral roots was counted. (n ≥ 20) ** = *P <* 0.01. (**B-C**) Arabidopsis DR5::GFP reporter lines were inoculated with the indicated bacterial strains for 48 hrs on agar plates. (B) 100x confocal images of GFP fluorescence from DR5::GFP expression. (**C**) Quantification of GFP fluorescence from maximum intensity projection images. Shown are average and SD n=5. *** = *P <* 0.005. Scale bar 50μm (**D**) Seedlings were inoculated with either WT or *ΔysnE* bacteria with or without 5μM IAA (for *ΔysnE* bacteria) for 48 hrs on agar plates and the number of colonizing bacteria was counted. Shown are averages and SD of 2 independent experiments (log_10_ transformed) with n ≥ 3 for each, *** = *P <* 0.005. (**E-F**) Seedlings were inoculated with either WT or *ΔysnE* bacteria expressing GFP (*amyE::Pspac-gfp*) for 48 hrs on agar plates. Shown are 200x confocal images of DIC from roots (left panels) and GFP fluorescence from bacteria (right panels) for WT (E) and *ΔysnE* (F) bacteria. Scale bars 50μm (**G-H**) seedlings were inoculated with either WT (G) or *ΔysnE* (H) Bacteria expressing GFP (*amyE::Pspac-gfp*) and followed by time lapse confocal microscopy for 12 hrs. Shown are 400x overlaid images of DIC from roots (grey) and GFP fluorescence from bacteria (green), taken at the indicated time points. Yellow arrows highlight bacterial growth arrest, probably culminating in cell death. Red arrows highlight *ΔysnE* bacteria replicating away from the root plane. Scale bars 10μm.

*ΔysnE* bacteria failed to colonize the root as efficiently as WT bacteria indicating that bacterial auxin not only triggers a root developmental response, but is necessary for efficient root colonization (Fig 1D-1F). Addition of exogenous auxin (IAA), restored *ΔysnE* bacterial colonization (Fig 1D). *ΔysnE* bacteria exhibited normal growth in vitro (Fig S1C), normal swarming motility (Fig S1D), and biofilm formation (Fig S1E), indicating that these processes, which can influence root colonization ^27, 28^, are not involved in the reduced colonization capacity. In an attempt to elucidate the underlying cause, we performed time lapse microscopy of root colonization by WT and *ΔysnE* GFP expressing bacteria ^20^. While most of the WT bacteria replicated and formed colonies over the root (Fig 1G), *ΔysnE* bacteria failed to replicate (Fig 1H). Most of the *ΔysnE* bacteria exhibited growth arrest or cell death (Fig 1H). Although some of the *ΔysnE* bacteria did replicate, their progeny failed to adhere to the root and spread away from it (Fig 1H). We conclude that bacterial auxin is necessary for *B. valezensis* to survive and replicate on the root.

### Auxin is necessary for *B. valzensis* to antagonize the plant immune reaction

The reduced ability of *ΔysnE* bacteria to colonize the root (Fig 1H) led us to hypothesize that *B. valzensis* is able to trigger a plant immune response. Auxin produced by pathogenic bacteria had previously been shown to reduce the plant immune response ^29^. RNA sequencing of whole roots after 48 hours of bacterial colonization (Table S1), revealed that gene categories related to immune system activation, like camalexin synthesis and callose deposition, were enriched in the root transcriptome as compared to buffer inoculated roots (mock) (Fig 2A and table S2). These early response results were corroborated by increased expression of immune related promoters (pPER5, pFRK1) fused to fluorescent reporters ^30^ (Fig S2A-S2B), as well as callose deposition (Fig S2C-S2D), indicating that *B. valzensis* colonization elicits an immune reaction. Monitoring plant ROS production also revealed a significant response (Fig 2B).

**Fig 2.**
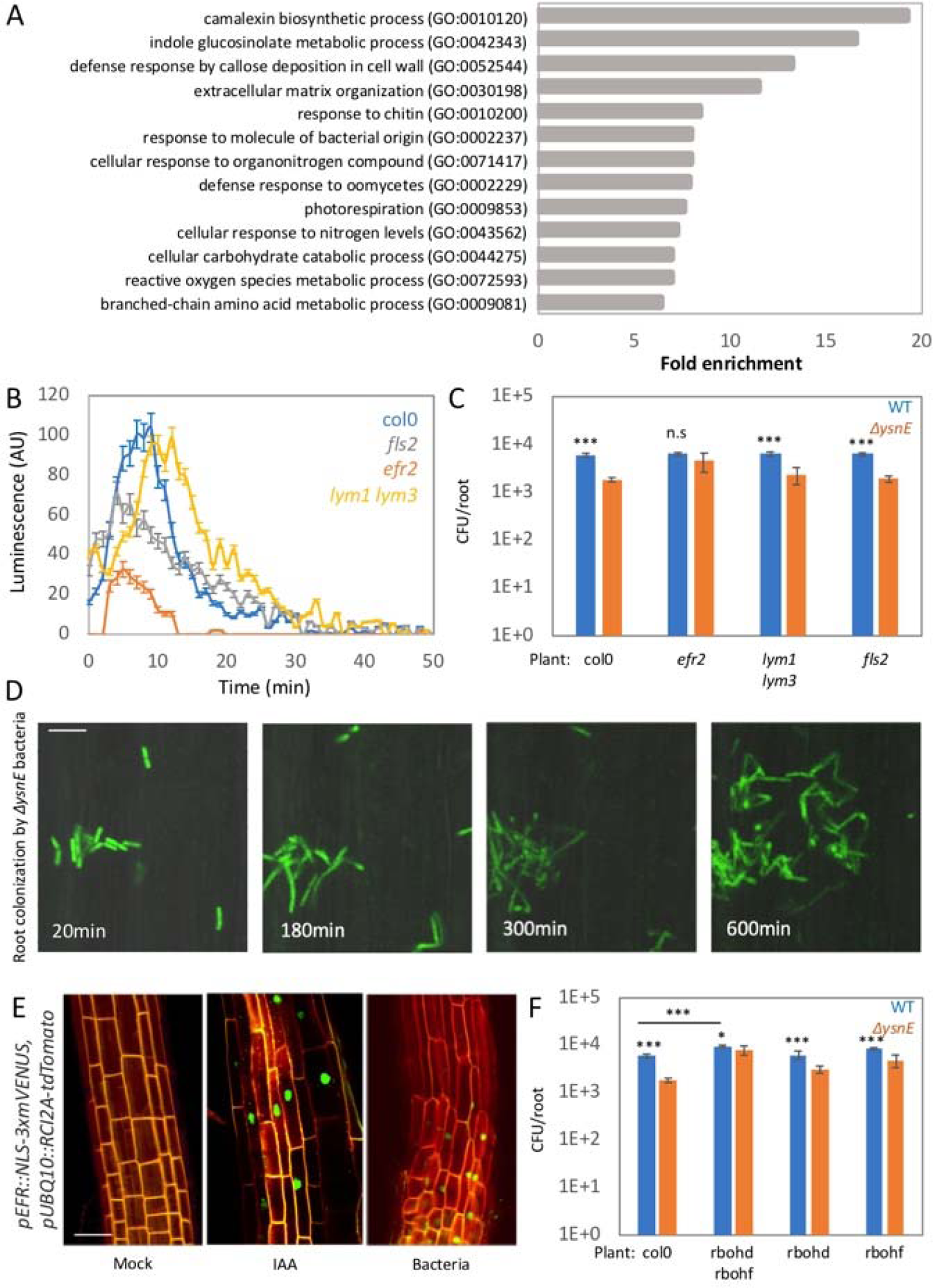
Bacterial auxin counteracts the plant immune response. (A) Representative biological process GO term analysis of plant genes upregulated in response to *B. valezensis* colonization. *P < 0.05* (see table S2 for the full list of GO categories). Leaf discs from 28 days old plant, taken from the indicated plant genotypes were incubated with bacteria adjusted to OD 0.1, and ROS burst was measured. Shown are average and SD (n ≥ 10). **(B)** Seedlings from the indicated genotypes were inoculated with either WT or *ΔysnE* bacteria for 48 hrs on agar plates and the number of colonizing bacteria was counted. Shown are averages and SD of 2 independent experiments (log_10_ transformed) with n=3 for each, *** = *P <* 0.005. Seedlings were inoculated with *ΔysnE* bacteria expressing GFP (*amyE::Pspac-gfp*) and followed by time lapse confocal microscopy for 12hrs. Shown are 400x overlay images of DIC from roots (grey) and GFP fluorescence from bacteria (green), taken at the indicated time points. *ΔysnE* bacteria replicated over the root, but failed to adhere as WT bacteria did (compare to figure 1G). Scale bar 10μm **(C)** Seedlings of *pEFR::NLS-3xmVENUS, pUBQ10::RCI2A-tdTomato,* were inoculated with WT bacteria, grown in the presence of 5μM IAA, or buffer for 48 hrs. Shown are 400x representative overlay images of *pUBQ10::RCI2A-tdTomato* (cell wall, red) and *pEFR::NLS-3xmVENUS* (EFR expressing cells, green) from 5 roots for each condition. Scale bar 20μm **(D)** Seedlings from the indicated genotypes were inoculated with either WT or *ΔysnE* bacteria for 48 hrs on agar plates, and the number of colonizing bacteria was counted. Shown are averages and SD of at least 2 independent replicates (log_10_ transformed),with n=3 for each. * = *P < 0.05*. *** = *P <* 0.005.

To identify the pathway by which the immune response is triggered, we measured the ROS response in plants deficient in three different MAMP receptors: *fls2,* mutant in the receptor for bacterial flagella ^31^*, efr2,* mutant in the receptor for bacterial elongation factor TU ^32^, and *lym1,lym3,* mutant in the receptor for bacterial peptidoglycan ^33^. We found that *efr2* exhibited the largest reduction in ROS production upon challenge with *B. valzensis* (Fig 2B). *efr2* also exhibited a reduction in callose deposition (Fig S2C-S2D). We hypothesized that if bacterial auxin is necessary to antagonize the plant immune response, then colonization of bacteria deficient in auxin production would be restored on mutant plants with compromised immunity. Consistent with this hypothesis, we found that *ΔysnE* growth was significantly enhanced on *efr2* mutant plants (Fig 2C-2D and Fig S2E). Although these auxin-deficient bacteria were able to replicate on *efr2* mutant plants, unlike WT bacteria they do not adhere to the root (compare Fig 2D and Fig 1G). The EFR2 receptor is expressed in the roots at very low levels (Fig 2E) ^30, 34^. However, *B. valzensis* colonization highly stimulated expression of a pEFR transcriptional reporter (Fig 2E). Intriguingly, exogenous IAA also stimulated pEFR reporter expression (Fig 2E), suggesting that bacterial auxin is also able to stimulate EFR expression. We conclude that intact bacterial auxin synthesis is necessary for the bacteria to survive the plant immune response.

### Auxin antagonizes ROS toxicity

To identify the components of plant immunity perturbed by bacterial auxin, we monitored *ΔysnE* bacterial colonization on mutants in immune response genes. Mutations in *crt3* essntial for EFR2 receptor function ^35^, restored *ΔysnE* bacterial colonization (Fig S3A), while neither perturbation in the indole-glucosinolate and camalexin synthesis pathway (*myb51* ^36^ and *cyp71a13* ^37^) nor defects in plant stress hormone effectors (*npr1-5*, *ein2-5*, *jar1-1*) affected *ΔysnE* bacterial colonization (Fig S3A). The lack of SA response [*npr1-5,* ^38^] is notable, as auxin is known to antagonistically interact with the SA pathway to enhance the colonization of *P. syringae* ^29, 39^. In contrast, the growth of auxin deficient bacteria was restored on *rbohd,rbohf* plants, which are defective in immune triggered ROS production (Fig 2F) ^40^. Moreover, significant recovery of *ΔysnE* bacterial colonization was obtained when ROS production was chemically inhibited by DPI ^41^ (Fig S3B). These results suggest that bacterial auxin antagonizes plant ROS production to enable root colonization.

On *rbohd,rbohf* plants, *B. valezensis* caused a pathogen-like effect with extensive overgrowth (Fig 2F), reduced numbers of lateral roots, and smaller plants (Fig S3C-S3D). This indicates that *efr2* plants, although perturbed in *B. valezensis* triggered immunity are still capable of eliciting a sufficiently strong immune response with ROS production (Fig 2B) to keep the bacteria from overgrowing the plant. ROS are toxic molecules, utilized by the plant to kill invading pathogens [e.g. ^42^].

NADPH oxidase enzymes like RbohD and RbohF produce superoxide (O^-^) ions, which can be further converted into other reactive oxygen species like H_2_O_2_ ^43^. O^-^ was highly toxic to *B. valzensis* in vitro (Fig 3A), while H_2_O_2_ killed bacteria only at a high concentration (500 μM) (Fig S3E). O^-^ was significantly less toxic to WT bacteria, in comparison to auxin deficient bacteria (Fig 3A). Exogenous IAA enhanced the survival of both bacteria (Fig 3A). These results suggest that auxin enables the bacteria to survive the toxic effects of ROS.

**Fig 3.**
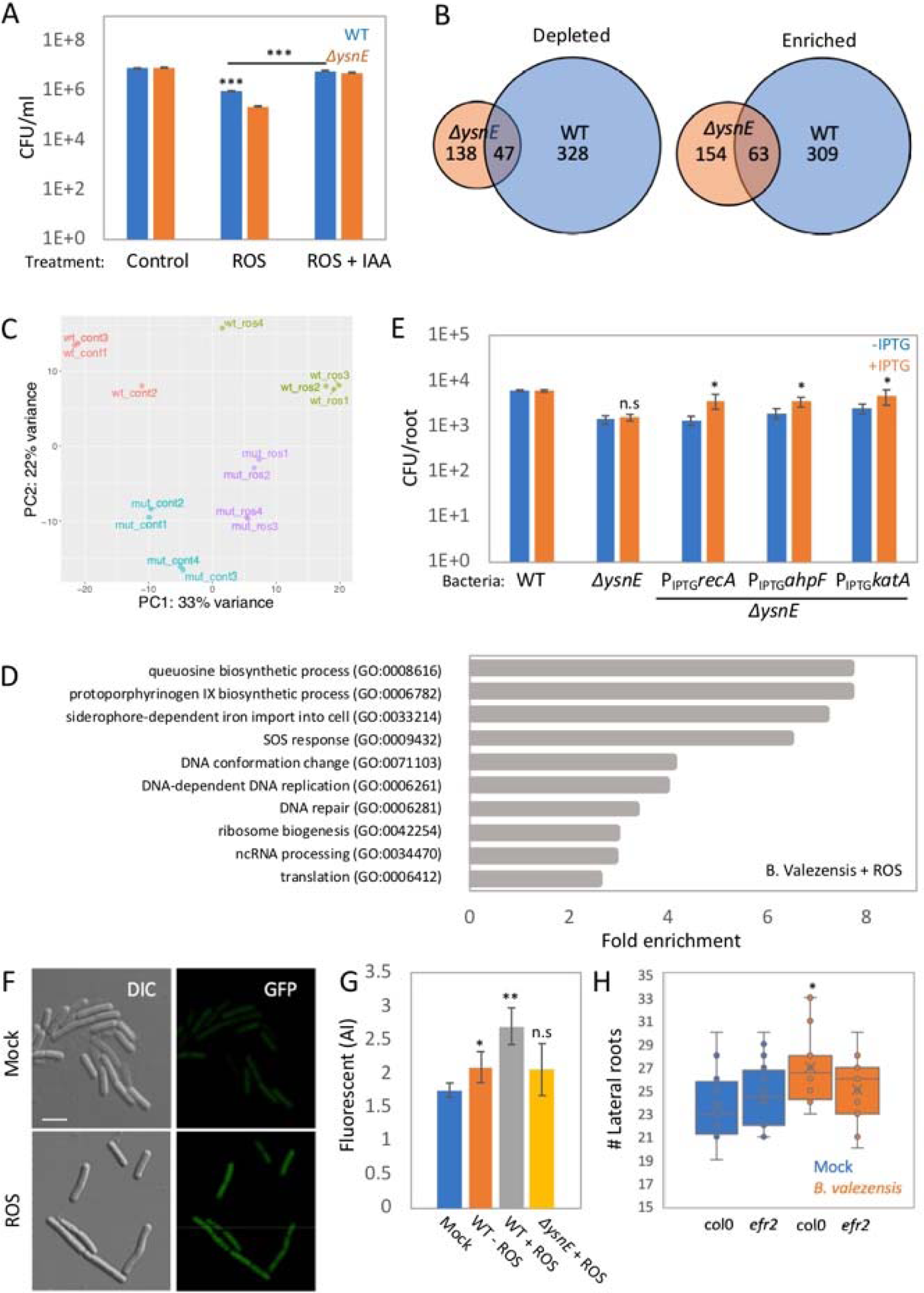
Bacterial auxin counteracts ROS toxicity. (A) Bacterial cultures grown to OD_600_=1 were treated with O^-^, in the presence or absence of 5μM IAA for 30min, and CFU were counted. Shown are averages and SD (log_10_ transformed) n=3. *** = *P <* 0.005. **(B)** Venn diagram illustrating significantly affected genes. On the left are genes depleted after O^-^ treatment, on the right genes enriched after O^-^ treatment. **(C)** Principal component analysis of RNA sequenced from WT or *ΔysnE* bacteria treated with O^-^ at sub-lethal concentration (see Fig S3F) for 30min, or with the substrate alone as control **(D)** Representative biological processes GO term analysis of genes upregulated in response to O^-^ treatment. *P < 0.05.* (see table S4 for the full list of GO categories) **(E)** Seedlings were inoculated with the indicated bacterial strains on plates containing 0.5mM IPTG (+IPTG) or in the absence of IPTG (-IPTG) for 48 hrs and the number of colonizing bacteria was counted. Shown are averages and SD of 2 independent experiments (log_10_ transformed), with n 3 for each. * = *P < 0.05.* **(F)** Ysne-GFP expressing bacteria grown to OD_600_=1 were treated with O^-^ for 30min and observed under the microscope. Shown are 400x DIC image (left) and GFP fluorescence from YsnE (right). Bacteria treated with the substrate without the enzyme (Mock) used as control. Scale bar 3μm. **(G)** Arabidopsis DR5::GFP reporter lines were inoculated for 12hrs with media derived from WT or *ΔysnE* bacteria grown to OD_600_=1 and treated with O^-^ for 30min. Shown are average and SD of fluorescent intensity from DR5::GFP, (n 5). ** = *P < 0.005*. * = *P < 0.05*. **(H)** Seedlings of Col-0 or *efr2* plants were inoculated with bacteria or buffer (mock) on agar plates for 7 days and the number of lateral roots was counted. (n≥ 20). * = *P < 0.05*.

To gain a deeper understanding of the effect of auxin on bacterial interaction with ROS, we examined global gene expression changes in WT and *ΔysnE* bacteria after addition of O^-^. In WT bacteria, 371 genes were upregulated and 374 genes downregulated (Fig 3B), while *ΔysnE* bacteria exhibited a weaker response (Fig 3B-3C), with only 153 genes upregulated and 184 downregulated (Fig 3B and table S3). Enriched GO categories for upregulated genes in WT bacteria included SOS response, and DNA repair, while the DNA repair category was missing in *ΔysnE* (Fig 3D and table S4). *katA* and *ahpF* genes are important for ROS detoxification ^44, 45^, and the *recA* gene is important for DNA repair ^46^. All three were upregulated in response to ROS treatment (table S3, all three were induced to a greater extent in WT bacteria.). Expression of these genes in *ΔysnE* bacteria, under an IPTG inducible promoter significantly enhanced root colonization (Fig 3E). Interestingly, GO categories related to iron homeostasis, were enriched in the transcriptome of WT bacteria but not in *ΔysnE* bacteria (Fig 3D and table S4). Ferrous (Fe^2+^) iron is known to interact with hydrogen peroxide in a Fenton reaction to produce a toxic hydroxyl radical ^47^, thus iron sequestration can protect cells from the toxic effects of ROS. Expression of the siderophore bacillibactin or heme synthesis operons in *ΔysnE* bacteria under IPTG inducible promoters enhanced their ability to colonize the root (Fig S4A). Lowering the iron content of the MS media by 50% also improved root colonization by *ΔysnE* bacteria (Fig S4B). Of note, auxin was able to protect *B. valezensis* from iron toxicity in vitro (Fig S4C-S4D). Among the significantly depleted gene categories in WT bacteria were TCA cycle, and carbohydrate and amino acid transport, though none of these categories was depleted in *ΔysnE* bacteria (Fig S4E and table S4). We speculate that WT bacteria enter a growth arrest that can protect them from ROS toxicity, while *ΔysnE* bacteria that fail to induce growth arrest are killed. Thus, our results establish ROS as a major limiting factor during root colonization and auxin as a key bacterial effector to mitigate ROS toxicity. Addition of IAA to bacteria without ROS had negligible effects on transcription (table S3), suggesting that auxin alone is not sufficient to explain these transcriptional changes and that other factors induced during stress are necessary for auxin to have its effect.

Given our findings that auxin plays a major role in mitigating ROS toxicity, we hypothesized that ROS exposure leads to auxin accumulation in bacteria. To test this hypothesis we fused YsnE to GFP and observed that it accumulated upon ROS treatment in-vitro (Fig 3F and Fig S5A). We collected the supernatant from bacterial cultures treated with ROS and applied it to DR5::GFP expressing plants, which led to a greater increase in DR5::GFP fluorescence as compared to plants treated with the supernatant from *ΔysnE* bacteria or from untreated WT bacteria (Fig 3G and Fig S5B). Consistent with these results, *efr2* roots colonized by bacteria failed to exhibit lateral root stimulation, and had normal primary root length (Fig 3H and Fig S5C-S5D), suggesting that EFR induced ROS production by the plant is necessary to trigger efficient bacterial auxin production.

### Auxin promotes bacterial adhesion and colony formation on the root

Although growth of *ΔysnE* bacteria is restored on *efr2* roots (Fig 2C-D and Fig S2E), these bacteria do not adhere to the root like WT bacteria does to Col-0 roots (Fig 2D). Interestingly, similar inefficient adhesion occurred when WT bacteria colonized *efr2* roots (Fig 4A-4B and Fig S6A-S6B). Quantification of root adhesion from time lapse microscopy revealed that, on average, 83% of bacteria colonizing WT Col-0 roots remain adhered to the root during a 12 hour experiment (Fig 4A) while only 32% did so on *efr2* roots (Fig 4A and Fig S6A). The macro-structure of bacteria colonizing a root after 48 hours revealed large clusters on Col-0 roots (Fig 4B and Fig S6B), while bacteria colonizing *efr2* roots were in small patches (Fig 4B and Fig S6B), probably reflecting the same phenomenon of perturbed adhesion and colony formation. This suggests that efficient ROS response, perturbed in *efr2* roots, is necessary for tight root adhesion and colony formation. Addition of exogenous IAA stimulated colonization on Col-0 as well as *efr2* plants (Fig 4C). Exogenous IAA can stimulate root colonization on mutant plants impaired in auxin perception (Fig S6C-S6D), suggesting that auxin, at least in part, affects bacterial ability to adhere to the root rather than the root’s response to bacteria.

**Fig 4.**
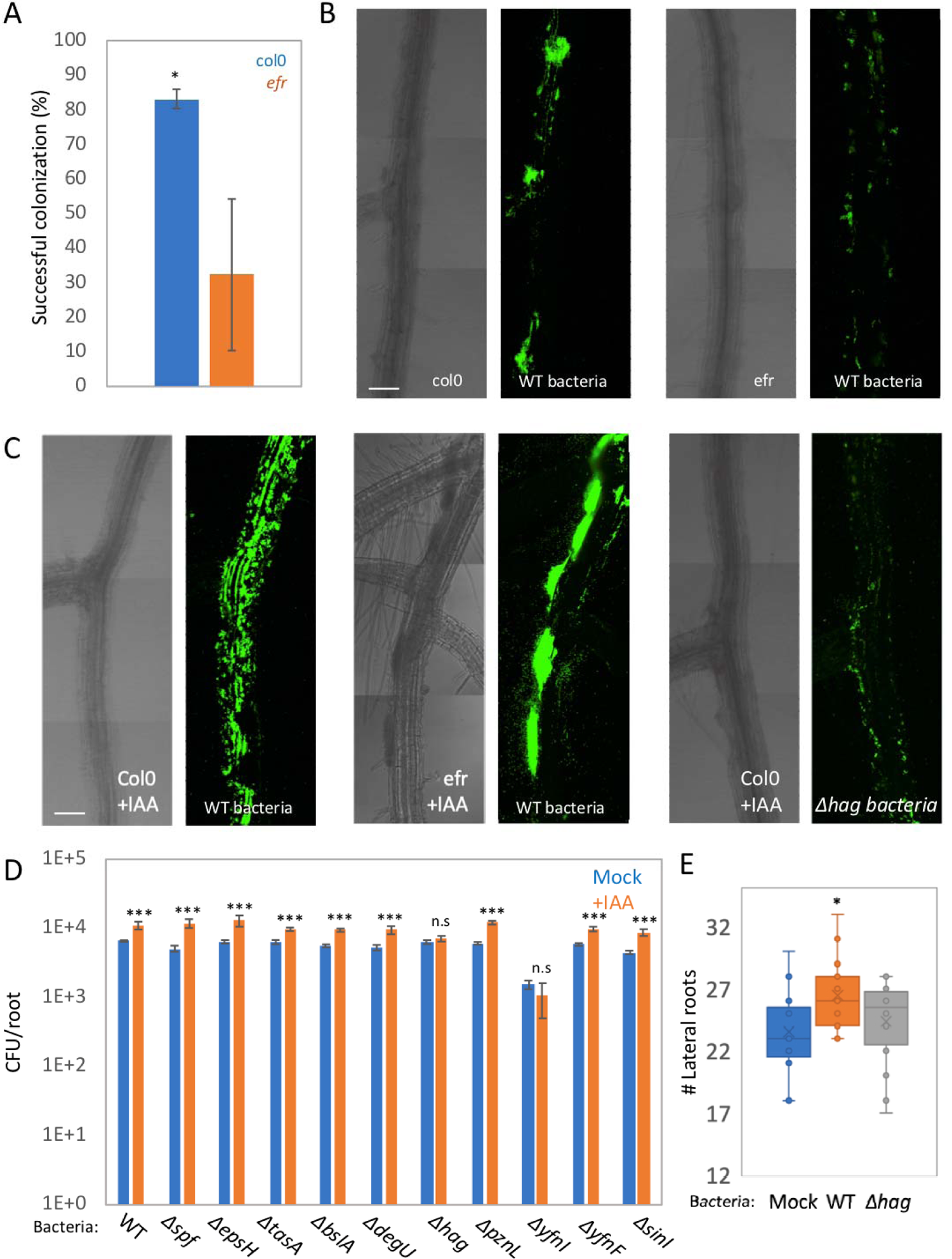
ROS induces auxin enhanced root colonization through stimulation of bacterial flagella. **(A)** Col-0 or *efr2* seedlings were inoculated with GFP expressing bacteria (*amyE::Pspac-gfp*) and followed by time lapse confocal microscopy for 12hrs. Spots of colonizing bacteria were counted at t = 2hr and followed until t =12hr. Bacteria remained attached during this time course were counted as successful colonization events (see an example in Fig S6A). Shown are percentages of successful colonization at t = 12 hrs of total spots of colonizing bacteria at t =2 hrs. The results are average and SD from at least 3 time-lapse experiments for each genotype. n≥ 25 spots for each time lapse. * = *P <* 0.05. (**B-C**) Seedlings were inoculated with the indicated bacterial strains expressing GFP, for 48 hrs on agar plates. In the absence (B) or presence of 5μM IAA (C). Shown are 200x confocal images of DIC from roots (left panels) and GFP fluorescence from bacteria (right panels). Scale bars 50μm (D) Seedlings were inoculated with the indicated bacterial strains with or without 5μM IAA for 48 hrs on agar plates and the number of colonizing bacteria was counted. Shown are averages and SD (log_10_ transformed), n = 3. *** = *P<* 0.005 Seedlings were inoculated with either WT or *Δhag* bacteria or buffer alone (mock) on agar plates for 7 days and the number of lateral roots was counted. n ≥ 20, * = *P <* 0.05.

To elucidate the mechanism by which auxin promotes root adhesion and spreading, we screened an array of colonization related mutant bacteria, impaired in motility, adhesion and biofilm formation genes ^48^ for auxin enhanced colonization (Fig 4D). Bacteria, with a mutated lipoteichoic-acid synthase gene, Δ*yfnI,* lost their ability to colonize the root, irrespective of IAA addition. Bacteria lacking a flagellar apparatus, Δ*hag,* colonized the roots similar to WT bacteria. However, they failed to exhibit enhanced colonization following IAA addition (Fig 4C-4D). Auxin induced flagellar formation was also suggested by the RNA sequencing analysis (table S3, IAA induced *hag* gene expression logFC=0.79) and in vitro motility assay (Fig S6E). Δ*hag* bacteria also failed to induce lateral root formation (Fig 4E). Thus, auxin induced flagella production is able to enhance root colonization necessary for lateral root stimulation.

### Plant immune system stimulates root colonization and auxin secretion by diverse bacterial species

To determine if bacterial auxin secretion and plant immunity interact in a similar manner for other bacteria we analyzed the colonization capacity of *Paenibacillus polymyxa* (*P. polymyxa*), a gram-positive bacteria known to secrete high amounts of auxin and stimulate plant growth ^49^. *P. polymyxa* stimulated lateral root formation and DR5::GFP expression in roots on agar plates (Fig 5A and Fig S7A). *P. polymyxa* also stimulated plant ROS production in an FLS2 dependent manner (Fig S7B). On *fls2* plants, *P. polymyxa* failed to stimulate lateral root production (Fig 5A) and had longer primary roots as compared to Col-0 (Fig 5B), despite bacteria reaching higher CFU on *fls2* plants (Fig 5C), suggesting that immune system activation and ROS production is necessary for bacterial auxin production. Finally, exogenous IAA further stimulated root colonization by *P. polymyxa* (Fig 5D). Thus, *P. polymyxa* produced auxin and plant immunity interact with each other, despite being recognized by a different immune receptor than *B. valzensis*. *Arthrobacter mf161* is another Gram-positive auxin secreting bacterium isolated from Arabidopsis roots ^50^. Inoculation by this bacterial strain stimulated lateral root formation and DR5::GFP expression (Fig S7C-S7D) as well as triggering the immune response in an FLS2 dependent manner (Fig S7E). *Arthrobacter mf161* failed to enhance lateral root formation, and had longer primary roots, on *fls2* plants (Fig S7C and Fig S7F). No difference in root colonization was observed between col-0 and *fls2* plants (Fig S7G) Finally, exogenous IAA further stimulated root colonization by this bacterial strain (Fig S7H). On the other hand, auxin didn’t stimulate root colonization of auxin secreting *Pseudomonas* species *wcs365* ^51^ *and wcs374* ^52^ (Fig S7I), suggesting that auxin stimulated colonization is not a general phenomenon but is bacterium-specific.

**Fig 5.**
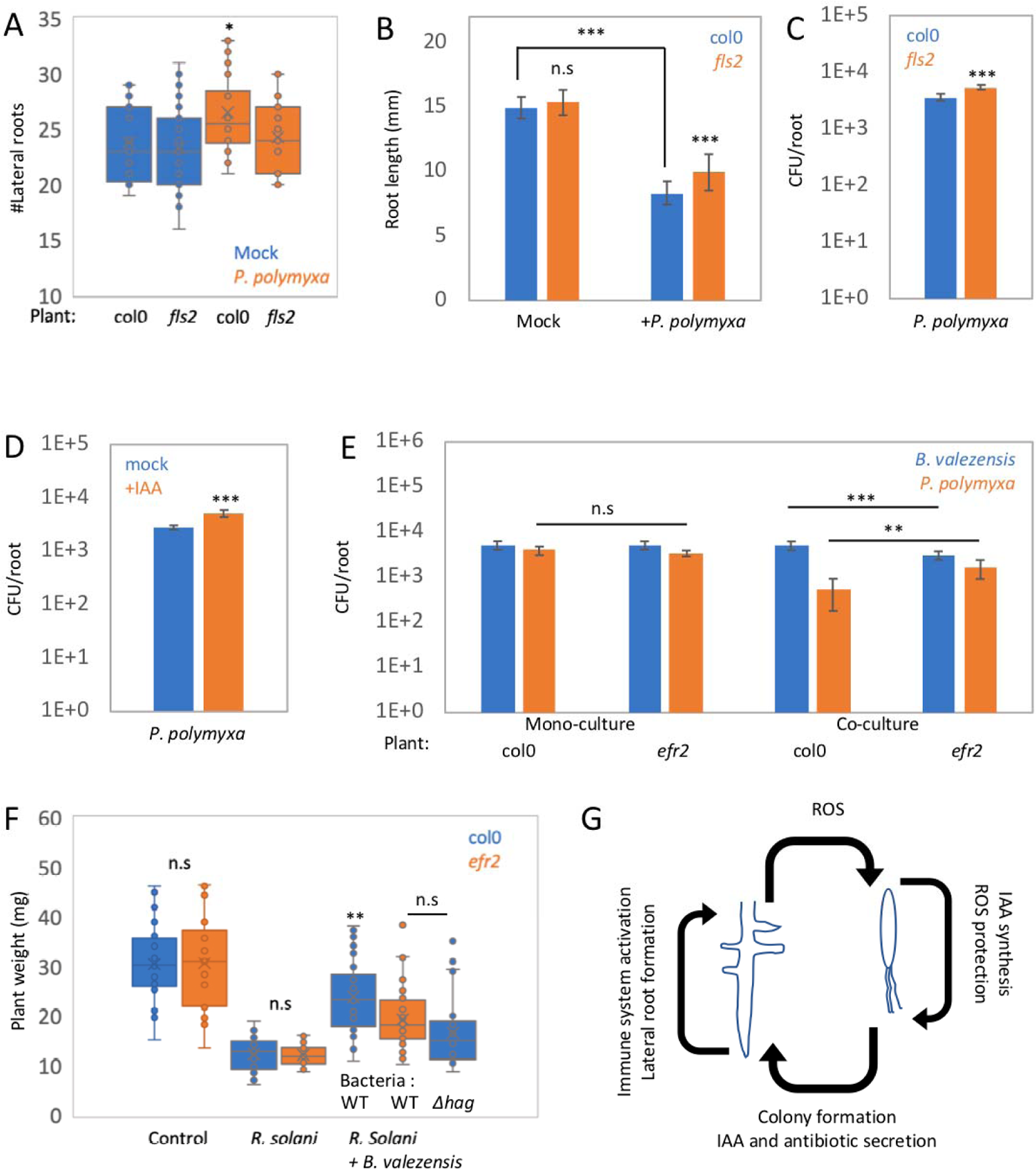
Plant immunity interaction with bacterial auxin secretion in *P. polymyxa*. (A) Col-0 or *fls2* seedlings were inoculated with *P. polymyxa* or buffer (mock) on agar plates for 7 days and the number of lateral roots was counted, n ≥ 20. * = *P <* 0.05. (B) Col-0 or *fls2* seedlings were inoculated with *P. polymyxa* or buffer alone (mock) on agar plates for 7 days and the length of the primary root was measured, n ≥20. *** = *P <* 0.005. (C) Col-0 or *fls2* seedlings were inoculated with *P. polymyxa* for 48 hrs on agar plates and the number of colonizing bacteria was counted. Shown is an average and SD of 2 independent replicates (log_10_ transformed), with n=3 for each. *** = *P < 0.005*. (D) Seedlings were inoculated with *P. polymyxa* for 48 hrs on agar plates with or without 5μM IAA and the number of colonizing bacteria was counted. Shown are averages and SD of 2 independent replicates (log_10_ transformed), with n=3 for each. *** = *P < 0.005*. (E) Seedlings were inoculated with either *P. polymyxa* or *B. valezensis* alone (monoculture) or in a mixture (1:1 ratio, co-culture) for 48 hrs on agar plates and the number of colonizing bacteria from each strain was counted. Shown are averages and SD of 2 independent replicates (log_10_ transformed), with n=3 for each. *** = *P < 0.005.* ** = *P < 0.01*. (D) Col-0 or *efr2* Seedlings were inoculated with WT or *Δhag* (only col-0) *B. valezensis,* or buffer for 48 hrs on agar plates. Then plates were inoculated with *R. solani* and incubated for an additional 7 days and plant weight was measured. Untreated plants (neither bacteria, nor fungi) were used as control. Shown are averages and SD, n ≥20. (E) Model describing the feedback loop between plant immune system activation and bacterial auxin secretion.

Our results indicate that ROS production by the plant immune system is necessary for efficient root adhesion. However, this phenomenon is not manifested in differences in bacterial load (Fig 4). Given that bacteria in nature compete with many other species to inhabit the same plant root niche ^1, 2^, we hypothesized that differences in *B. valezensis* root adhesion ability would become evident during competition with other bacteria. To test this hypothesis, we co-inoculated *P. polymyxa* and *B. valezensis* on Col-0 and *efr2* plants. *P. polymyxa* is only recognized by the FLS2 receptor but not by EFR2 (Fig S7B), while *B. valezensis* is mainly recognized by the EFR2 receptor (Fig 2B). After co-inoculation, *B. valezensis* outcompeted *P. polymyxa* on Col-0 (Fig 5E). However, on *efr2* plants, we observed a significant increase in *P. polymyxa,* concomitant with a reduction of *B. valezensis* colonization (Fig 5E). Co-inoculation of *B. valezensis* and *Arthrobacter mf161* on Col-0 and *efr2* plants had no significant effect on either bacteria (Fig S8A). Inspection of colonization sites revealed that *B. valezensis* and *P. polymyxa* heavily colonize the elongation and maturation zones of the root (Fig S8B_1-2_) while *Arthrobacter mf161* is largely absent from these regions and colonizes differentiated parts of the root (Fig S8B_3-4_). Our results suggest that immune system enhanced colonization affects *B. valezensis* and *P. polymyxa* competition as both compete for the same niche but not *B. valezensis* and *Arthrobacter mf161* competition, as they colonize different niches.

*B. valezensis* produces secondary metabolites that can inhibit the growth of plant fungal pathogens ^15^. We asked if plant immune system activation, triggering *B. valezensis* colony formation, enhances its ability to inhibit plant pathogen infection. We colonized Col-0 and *efr2* plants with *B. valezensis*, and infected the plants with the fungal pathogen *Rhizoctonia solani* ^53^.

*B. valezensis* inhibits the growth of *R. solani* in vitro (Fig S9A) and is able to protect the plants from fungal infection (Fig S9B) ^19^. Of note, plant protection was significantly better on Col-0 plants, as measured by plant weight, although *efr2* has no effect on fungal infection per se, (Fig 5F *R. solani* alone). EFR2 activation modulates *B. valezensis* colony formation, but may also enhance plant survival through induced systemic resistance (ISR) ^54^. To further differentiate between these effects, we measured plant protection by Δ*hag B. valezensis* (Fig 5F). Δ*hag* bacteria failed to protect the seedlings from *R. solani* infection (Fig 5F), despite inducing an immune response in the plant, similar to WT bacteria (Fig S9C). Thus, we conclude that the enhanced colony formation of *B. valezensis* on immune competent plants enables it to better protect the plant from fungal infection.

## Discussion

Our results are consistent with the presence of a feedback loop between the plant immune system and bacterial auxin secretion (Fig 5G). Root colonization by bacteria triggers an immune response and ROS production. ROS, in turn, elicits bacterial auxin production to mitigate ROS toxicity. Auxin promotes bacterial spreading over the root, and colony formation, while also inducing the expression of plant immune receptors, further accelerating the feedback loop. This enhanced colonization promotes the ability of *B. valezensis* to inhibit plant pathogenic fungus. Thus, a feedback loop between bacteria and the plant immune system promotes the fitness of both partners.

Recent work has elucidated the role of the plant immune system in shaping the normal root microbiota, in addition to fighting pathogens ^11, 12^. In these studies, an immune reaction was viewed as a negative factor for root colonization, shaping the microbiota by preventing bacterial overgrowth. Consistent with this view, our results show that *B. valezensis* overgrow on *rbohd/ rbohf* plants completely lacking ROS production (Fig 2F). However, bacteria grow on *efr2* plants, with partially perturbed immunity demonstrating that plant immune system activation also plays a positive role for bacterial colonization, triggering induction of auxin production by bacteria necessary for efficient root adhesion and colony formation.

Our results suggest that immune system activation interacts with bacterial auxin secretion to enhance bacterial colonization irrespective of the specific immune receptor, as we provide evidence that a similar feedback loop exists during *P. polymyxa* and *Arthrobacter mf161* colonization, despite being recognized by the FLS2 receptor rather than the EFR2 receptor. Thus, we uncovered a novel aspect of bacterial interaction with the immune system.

A prevalent view of mutualistic interactions is that symbiosis evolved through exploitative interactions that became attenuated over evolutionary time ^55–57^. Parallels were found between the immune system signaling pathway and the symbiotic association between plants and specialized mutualists, like the interaction between legumes and rhizobia ^55, 58^ as well as the association between plants and arbuscular mycorrhizal fungi ^59^. Our results reveal a more widespread relationship between plant immunity and beneficial bacteria, including non-specialized auxin secreting beneficial bacteria, potentially representing an earlier stage of the evolution of mutualism. Auxin is a key plant hormone, playing a wide range of roles in plant development ^60^.

Many bacterial species including pathogens like *Agrobacterium tumefaciens* and *Pseudomonas syringae* as well as beneficial bacteria like *Azospirillum brasilense* are known to synthesize auxin, and manipulate the plant through auxin secretion ^23, 61, 62^. However, despite decades of research on bacterial auxin production and how it affects plants, the role played by auxin on bacterial physiology is poorly understood. Previous studies found a bacterial transcriptional effect for auxin, but only at concentrations far above those that modify plant physiology ^63–65^ Our results suggest that auxin primarily affects the producer bacteria, acting as a stress related signal to protect them from reactive oxygen species. Mutations in the auxin synthesis pathway lead to profound transcriptional effects following ROS treatment. However, we failed to observe a substantial role for exogenous IAA under non-stressed conditions. This suggests that auxin may not be sufficient by itself to induce a significant response in bacteria similar to its effect on plants. Rather, auxin needs other factors that are induced during stress to have its effect. Further research will be necessary to elucidate the role played by auxin in bacterial physiology and stress adaptation.

Plants interact with a wide variety of bacterial species in nature. The composition of the plant microbiome is affected by factors such as soil geochemistry, bacterial diversity, the amount and composition of exudates, immune system activation, and by bacterial interaction with other bacteria, with phages or with other organisms. Understanding the effect of each of these components will enable rational manipulation of the plant microbiome to the benefit of the plant. Bacterial auxin production is highly prevalent among root colonizing bacteria ^66^, and the effect of auxin secreting and degrading bacteria in the root microbiome on plant physiology was recently explored ^67^. Here we have shown that auxin secreting bacteria interact with the plant immune system to promote their association with the plant and their competition with other bacteria.

## Acknowledgements

We thank G. Wachsman for help in RNA sequence analysis. We are greatful to S.Y He (Duke), X. Dong (Duke), B. Kunkel (WasU), and T. Nolan, R. Shahan, C. Winter from the Benfey lab, for critical reading of the manuscript. We thank J. Dangl (UNC Chapell Hill) and D. Russ from the Dangl lab for providing bacterial strains and for valuable discussions.

## Author contributions

ET and PNB conceived the project. ET performed the experiments. ET wrote an initial draft. PNB reviewed and edited the manuscript.

## Supplementary Materials

## Materials and methods

### Bacterial and fungal strains and growth conditions

WT *B. valezensis fzb42* bacteria and its mutant derivatives *ΔysnE*, *Δspf* and *Paenibacillus polymyxa ATCC842* were purchased from the Bacillus genetic stock center (http://www.bgsc.org/). *Arthrobacter mf161* was kindly provided by Prof. Jeff Dangl (University of North Carolina). *B. valezensis amyE::pSpac-GFP* was purchased from NORDREET company (https://www.nordreet.de/). Other *B. valezensis* mutant strains including: ΔepsH, ΔtasA, ΔbslA, ΔdegU, Δhag, ΔpznL, ΔyfnI, ΔyfnF, ΔsinI. IPTG inducible genes including: P*_IPTG_recA,* P*_IPTG_katA,* P*_IPTG_ahpF,* P*_IPTG_hemA,* P*_IPTG_dhbA,* and y*snE-gfp* were generated in this study. The media and growth conditions used for DNA transformation of *B. valezensis* were described in ^18^.

Gene deletions were performed by PCR amplification of 1000bp upstream and downstream of a given gene, the gene flanking regions were fused to an antibiotic resistance cassete using NEB builder (NEB) according to manufacturer’s instructions. The reactions were amplified by PCR for 30 cycles and transformed into *B. valezensis*. Bacteria with IPTG inducible genes were generated by PCR amplification of 1000bp on either side of a gene and cloned with an antibiotic resistance cassette, pHyperSpac promotor [from pdr111 plasmid Pdr111 and antibiotic resistance cassete kindly provided by Prof. David Rudner (Harvard)] and the gene. The 4 fragments were fused together using NEB builder (NEB). The reaction was amplified by PCR for 30 cycles and transformed into *B. valezensis*. y*snE-gfp* bacteria were generated by PCR amplification of *ysnE* without a stop codon, the GFP coding region from AR16 ^69^, an antibiotic resistance cassette, and 1000bp downstream of *ysnE*. The 4 fragments were fused together and transformed into *B. valezensis.* The primers used in this study are listed in Table s5. The bacteria were cultivated routinely on Luria broth (LB) medium. When needed the medium was solidified with 1.5% agar. For biofilm formation, bacteria were inoculated into MSgg medium and incubated without shaking for 4 days at 25° as described in ^70^. For experiments with IPTG inducible promoters (Fig 3B, Fig S4B), 0.5mM IPTG was added to the growth media 30min before root inoculation, and later bacteria inoculated onto roots, on plates containing 0.5mM IPTG. For O^-^ treatment, bacteria were grown to OD_600_ = 1, then 0.5mM xanthine added and 5μl xanthine oxidase enzyme (Sigma) (Fig 3A, fig 3H, Fig S5A) or 0.5μl enzyme for the RNA sequencing experiments. *Rhizoctoinia solani* isolate was kindly provided by Prof. Marc Cubeta (NCSU). Fungi were routinely grown on PDA plates (Sigma).

### Plant strains and growth conditions

The Arabidopsis (Arabidopsis thaliana) SALK, SAIL and CS series of transfer DNA insertion lines of *ein2-5* (CS3071), *npr1-5* (CS3724), *fls2*_SAIL, *efr2* (CS2103705), *lym1 lym3* (CS2103242), *rbohd* (CS68747), *rbohf* (CS68748), *rbohd rbohf* (CS68522), *tir1-1 afb4-8 afb2-3* (CS69646), *jar1-1 axr1-3* (CS67934), *myb51* (CS421816), *axr5-1* (CS16234), *cyp71a13* (CS879462), and *crt3* (CS2103723) mutant alleles were purchased from the Arabidopsis Biological Resource Center (http://www.arabidopsis.org/). *pFRK1::NLS-3xmVENUS pUBQ10::RCI2A-tdTomato*, *pPER5::NLS-3xmVENUS pUBQ10::RCI2A-tdTomato,* and *pEFR::NLS-3xmVENUS pUBQ10::RCI2A-tdTomato,* were kindly provided by Prof. Niko Geldner (University of Lausanne). All plants were grown on 0.5 MS media containing 1.1 gr Murashige and Skoog basal salts (in 500 ml ddH2O), 1% sucrose, 1% agar and 5 ml (in 500 ml ddH2O) MES (50 gr/l, pH=5.8 with NaOH). Plants were stratified for 2 d in a 4°C dark room and grown vertically for 4-10 days under long-day light conditions.

### Monitoring bacterial growth on plant roots

Bacteria from fresh colonies were grown in LB medium to an OD_600_ = 1.0 and then diluted 1:100 in PBSx1 for CFU measurements and microscopy, or 1:10^3^ for lateral roots and primary root measurements, yielding approximately 1×10^6^, or 1×10^5^ cfu/ml respectively. Six-day old seedlings were transferred onto square Petri dishes containing 0.5 MS but without sucrose. 2μL of bacterial dilution were put right above the root tip and left to dry for 2 min. The square plates were kept in a vertical position during the incubation time at 22°C under long-day light conditions (16 h light/8 h darkness) in a plant growth chamber. For bacterial CFU counting and microscopy, plants were incubated with bacteria for 48hrs. Then the inoculated plant roots were cut and washed three times in sterile water. For CFU counting the seedlings were transferred to a tube with 1 ml of PBSx1 and vortexed vigorously for 20 seconds, then the serial dilution was plated on LB plates. Measuring callose deposition was done as described in ^71^. For fungal infection, 6 day old seedlings grown on 0.5MS plates were inoculated with 10^-3^ CFU/ml of *B. valezensis* or buffer for 48 hrs, then a 5mm mycelial plug from the fungal culture was placed on the bottom of the plate and allowed to spread for an additional 7 days, after which, plant weight was measured.

### Microscopy

Roots were observed using a Zeiss LSM 880 laser scanning confocal microscope with the indicated lenses. Lateral root number was counted under a Zeiss Axio Zoom,V16 fluorescence dissecting scope at 10× magnification. Fluorescent intensity and length measurement were done using ImageJ.

### Measurement of plant ROS production

Leaf discs were cut with a 4 mm biopsy punch from 4 week-old plants and placed on sterile water with their adaxial side up in a white 96-well microtiter plate (Costar, Fisher Scientific) containing 150 μl H_2_O and then incubated overnight at 22°C in continuous light for 20 to 24 hours to reduce the wounding response. Immediately prior to elicitation, H_2_O was removed from each well and 100 μl of the elicitation solution (100μg/ml HRP (sigma), 1μM luminol (sigma) and bacteria adjusted to OD_600_=1) were added. Elicitation solution without bacteria was used as a control. Plates were analyzed every 1 min for a period of 45 min using a TECAN Infinite 200 PRO microplate reader with signal integration time of 0.5s.

### RNA extraction library preparation and computational analysis

For plant RNA, plant roots were cut and immediately frozen in liquid nitrogen. RNA prepared using RNeasy Plus Mini Kit (Qiagen) according to the manufacturer’s instructions. RNA-seq libraries were prepared using QuantSeq 3’ mRNA-Seq Library Prep Kit (Lexogen) according to the manufacturer’s instructions. Illumina NextSeq 500 High-Output 75bp single reads were aligned to the *Arabidopsis thaliana* genome, and differentially expressed genes analyzed on the BlueBee platform (https://www.bluebee.com/lexogen) with default parameters. GO annotation was analyzed on (http://geneontology.org/) with default parameters.

For bacterial RNA preparation, bacteria treated with O^-^ for 30 min were precipitated and bacterial pellets immediately frozen in liquid nitrogen. Pellets were then resuspended in 500μl lysis buffer (30LmM Tris, 10LmM EDTA, 10Lmg/mL lysozyme) for 30 min in 37°. RNA was prepared using the RNAzol reagent according to the manufacturer’s instructions. rRNA was removed using NEBNext® rRNA Depletion Kit (Bacteria) according to the manufacturer’s instructions. RNA-seq libraries were prepared using KAPA RNA HyperPrep Kit (Roche) according to the manufacturer’s instructions.

Illumina MiSeq v2 150bp PE reads were aligned to *B. valezensis fzb42* using Kallisto ^72^. Differentially expressed genes with logFC=0.5 and p-value < 0.01 were identified using the edgeR package. The full code was described in ^73^. Genes were annotated based on homology to the genome of *B. subtillis* 168, and GO annotation analyzed on (http://geneontology.org/) with default parameters with *B. subtillis* 168 based annotation. At least 72% of the differentially expressed genes from each comparision had homologs in the *B. subtillis* 168 genome.

## Supplementary figures

**Fig S1.**
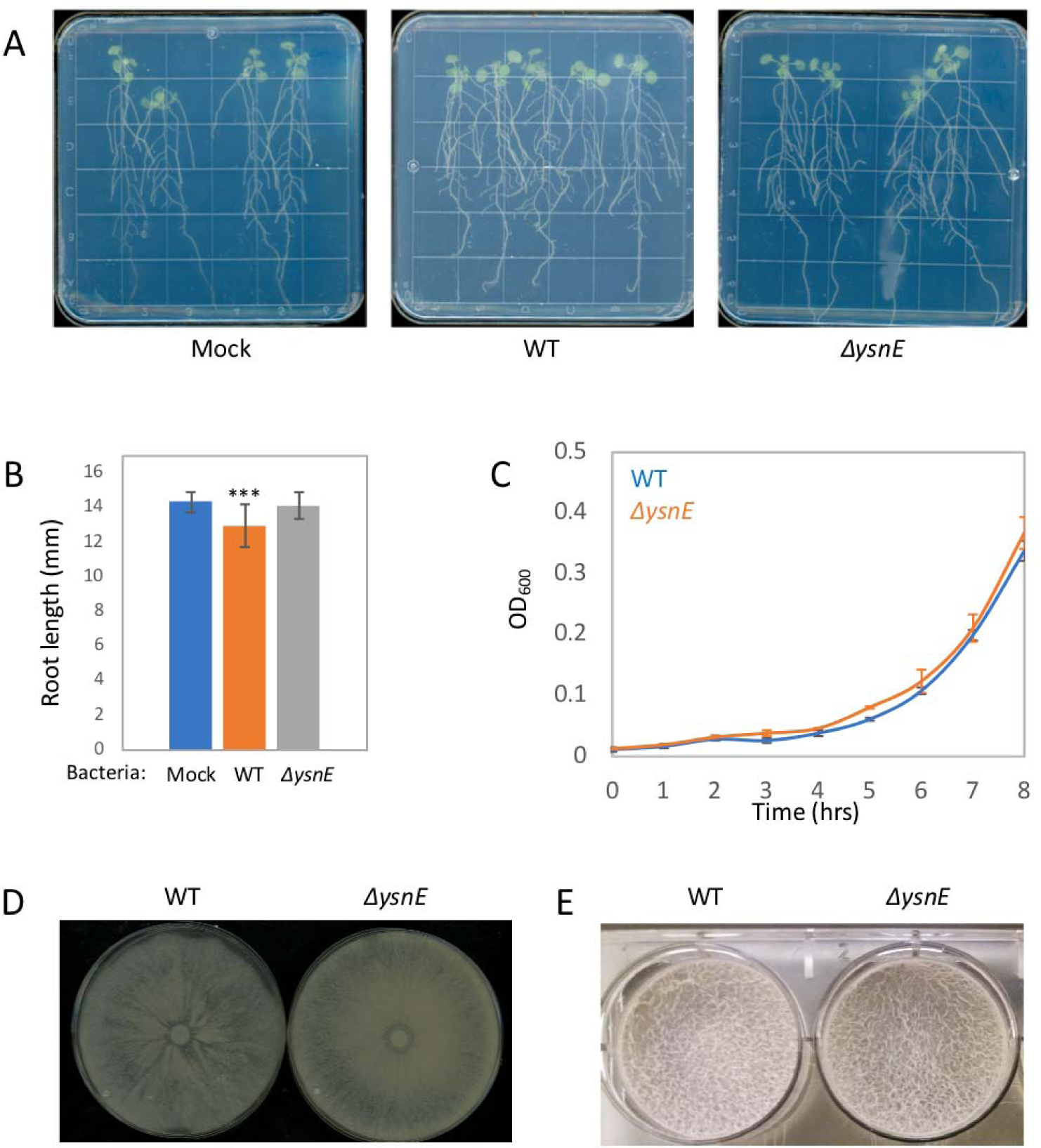
Bacterial auxin effects on the plant roots and bacterial physiology. (A) Seedlings were inoculated with either WT, *ΔysnE* bacteria, or buffer (mock) on agar plates for 7 days. Shown are representative plates from each of the conditions tested. (B) Seedlings were inoculated with either WT, *ΔysnE* bacteria or buffer alone (mock) on agar plates for 7 days and the length of the primary root was measured, n 20. *** = *P <* 0.005. (C) WT or *ΔysnE* bacteria were grown at 23° and OD_600_ measured. Shown are averages and SD, n = 3. (D) WT or *ΔysnE* bacteria were inoculated in the middle of 0.7% agar plates and incubated at 37° for 18 hrs. Shown are representative plates from 3 plates for each genotype. (E) WT or *ΔysnE* bacteria were inoculated into MSgg medium in 6 well plates and incubated at 23° for 96 hrs. Shown are representative wells from 3 for each genotype.

**Fig S2.**
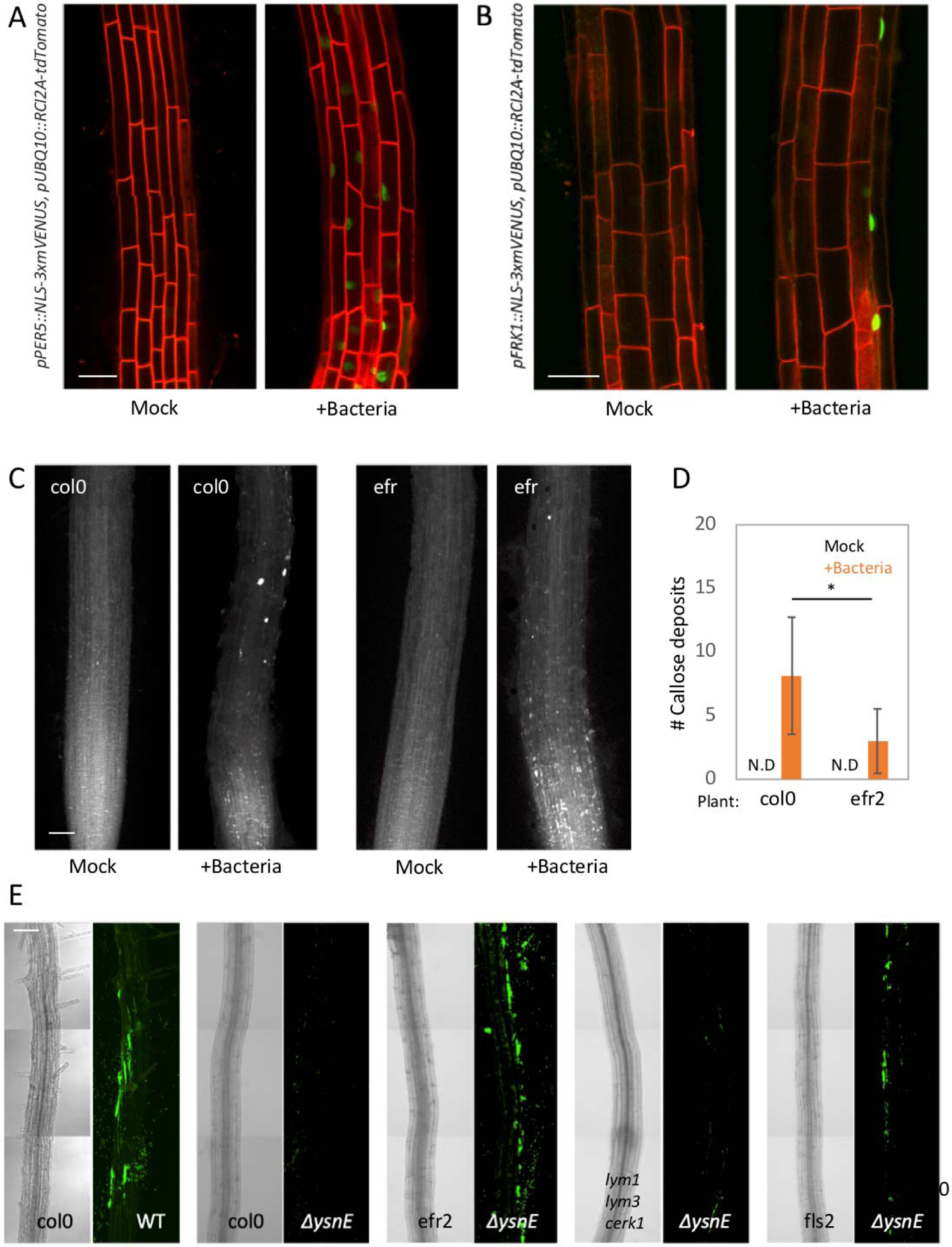
Bacteria induced an immune response in the root, in an EFR dependent manner. (A-B) Seedlings of the indicated genotypes, were inoculated with bacteria or buffer alone (mock) for 48 hrs. Shown are 400x overlay images of *pUBQ10::RCI2A-tdTomato* (red) and *pPER5::NLS-3xmVENUS* (green) (A), or *pFRK1::NLS-3xmVENUS* (green) (B). Representative roots from 5 roots from each condition. Scale bars 25μm (**C-D**) Seedlings of Col-0 or *efr2* were inoculated with bacteria or buffer alone (mock) for 48 hrs, then cleared with ethanol overnight and stained with aniline-blue dye. Shown are 200x confocal DAPI images (C) and average and SD from quantification of the number of callose deposits (D). n 4. * = *P <* 0.05. Scale bar 25μm (**E**) Seedlings from the indicated genotypes were inoculated with the indicated bacterial strains expressing GFP for 48 hrs on agar plates. Shown are 200x confocal images from roots (left panels) and GFP fluorescence from bacteria (right panels). Scale bar 50μm

**Fig S3.**
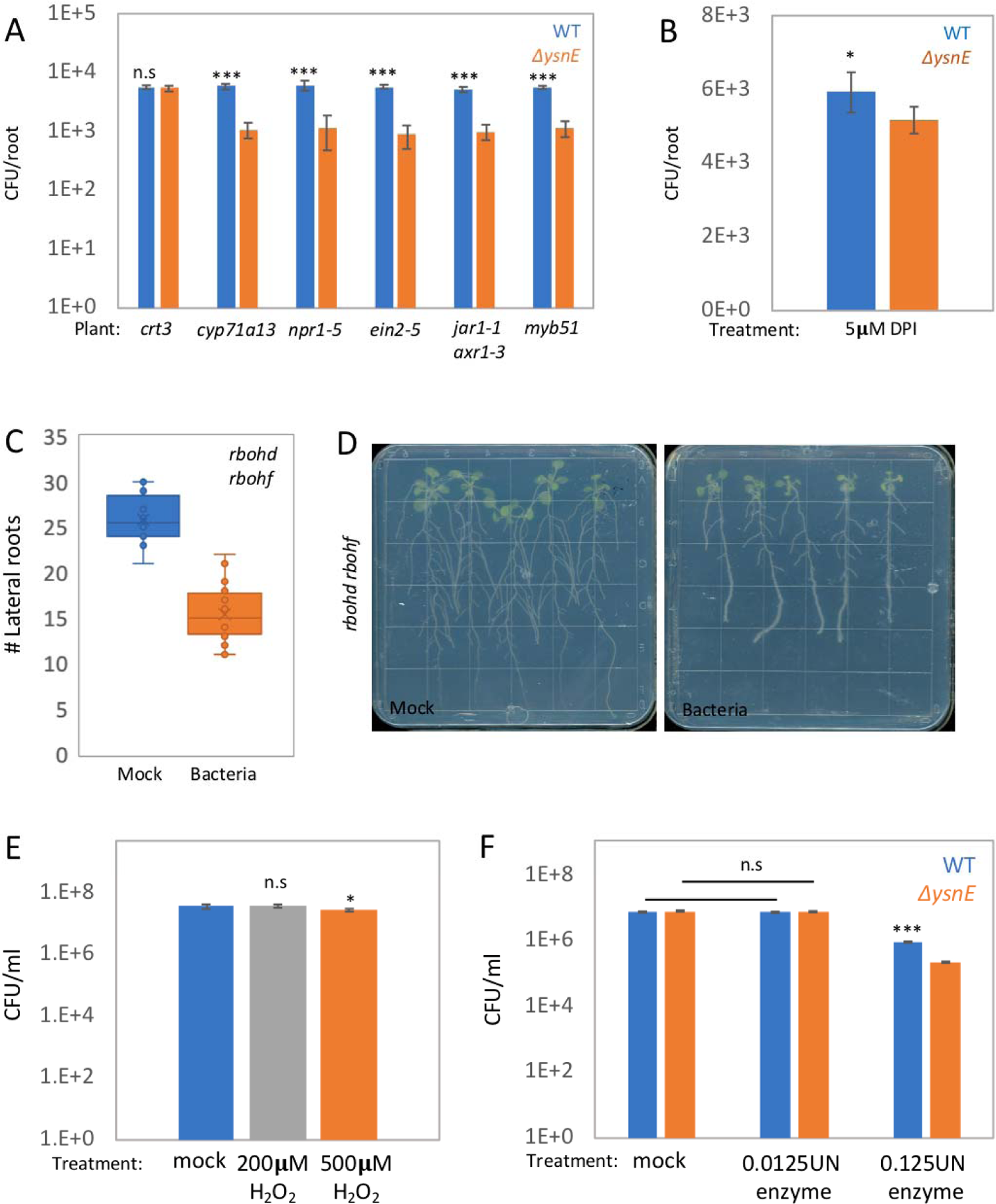
Bacterial auxin increases bacterial survival of plant immune system response. (**A**) Seedlings of the indicated genotypes were inoculated with WT or *ΔysnE* bacteria for 48 hrs on agar plates and the number of colonizing bacteria was counted. Shown are averages and SD (log_10_ transformed), n 3. *** = *P <*0.005 (B) Seedlings were inoculated with WT or *ΔysnE* for 48 hrs on agar plates containing 5μM DPI and the number of colonizing bacteria was counted. Shown are average and SD of 2 independent experiments, n = 3. * = *P <* 0.05. (C) Seedlings of *rbohd rbohf* plants were inoculated with bacteria or buffer (mock) on agar plates for 7 days and the number of lateral roots was counted, (n ≥ 20). (D) Seedlings of *rbohd rbohf* plants were inoculated with bacteria or buffer (mock) on agar plates for 7 days. Shown are representative plates from each of the conditions tested. (E) Bacterial culture grown to OD_600_=1 were treated with the indicated concentrations of H_2_O_2_ for 30min and CFU were counted. Shown are averages and SD (log_10_ transformed), n=3. * = *P < 0.05*. (F) Bacterial cultures grown to OD_600_=1 were treated with O^-^ through enzymatic reaction with xanthine and either 5μL xanthine oxidase enzyme (similar to Fig 3A) or 0.5μL xanthine oxidase enzyme (the concentration used for the RNA sequencing experiments, Fig 3C) for 30min and CFU were counted. Shown are averages and SD (log_10_ transformed), n=3. *** = *P < 0.005*

**Fig S4.**
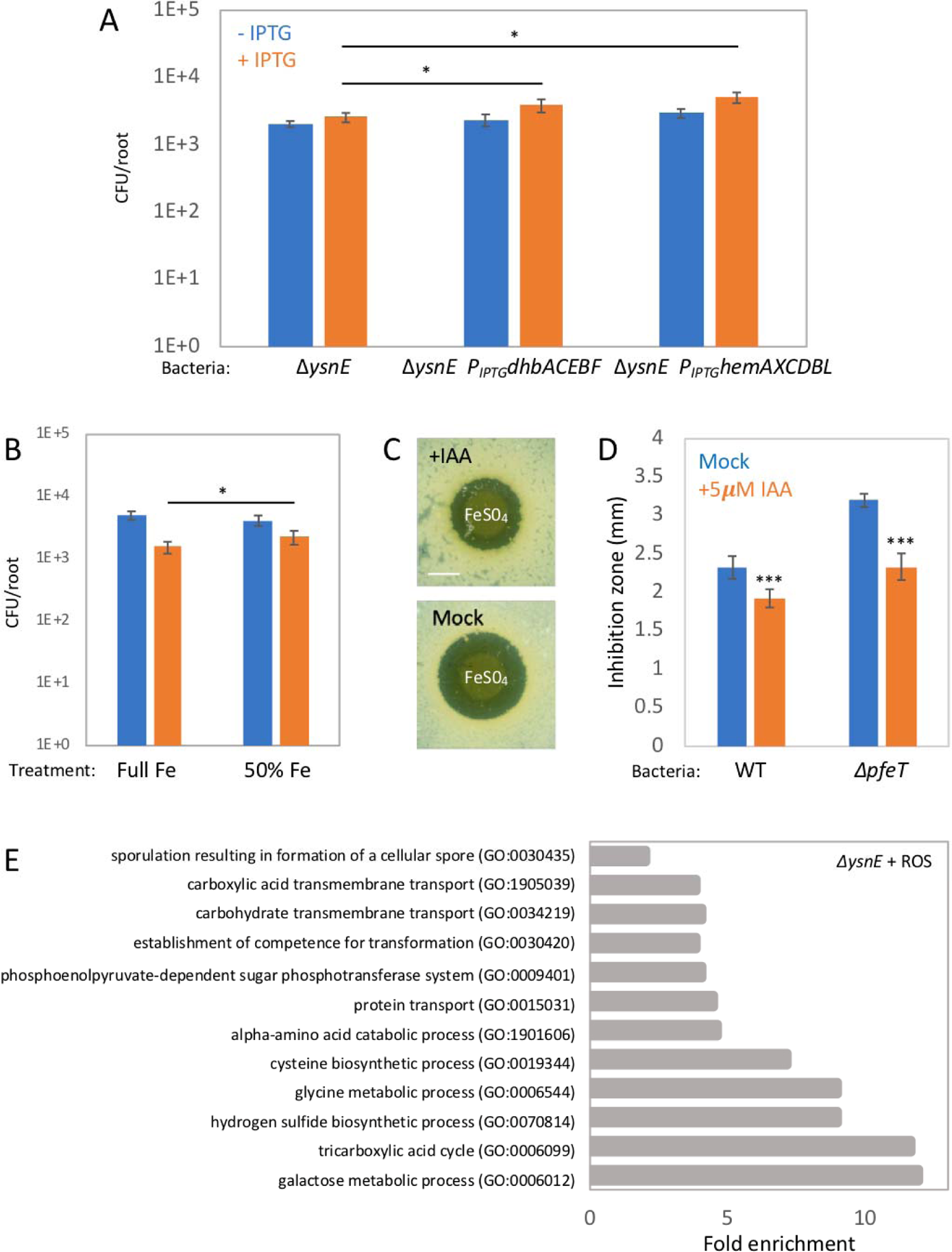
ROS induced genes are important for host survival of plant immune system response. (A) Seedlings were inoculated with the indicated bacterial strains on plates containing 0.5mM IPTG (+IPTG) or in the absence of IPTG (-IPTG) for 48 hrs and bacterial CFU counted. Shown are averages and SD of 2 independent experiments (log_10_ transformed) with n ≥ 3 in each. * = *P <* 0.05. (B) Seedlings were inoculated with WT or *ΔysnE* bacteria for 48 hrs on 0.5 MS agar plates without iron, supplemented with either 27.8 mg/l ferrous sulfate (full Fe) or 50% of that amount (half Fe) and the number of colonizing bacteria was counted. Shown are averages and SD (log_10_ transformed), n ≥ 3. * = *P <* 0.05. (**C-D**) The indicated bacterial strains were spread on LB agar plates with or without 5μM IAA. 5μL from 1M FeSO_4_ was spotted over the bacterial lawn, plates were incubated overnight at 37° and the growth inhibition zone around the spot was measured. Shown are images from ΔpfeT bacteria (D), which are mutant in the iron efflux pump ^68^. Grown in the presence (upper) or absence (lower) of 5μM IAA. Average and SD from quantification of growth inhibition zone of at least 6 FeSO_4_ spots for each treatment. *\*\*\* = P <* 0.005. Scale bar 0.5mm (**E**) Representative biological process GO term analysis of the genes depleted in response to O^-^ treatment. *P < 0.05*.

**Fig S5.**
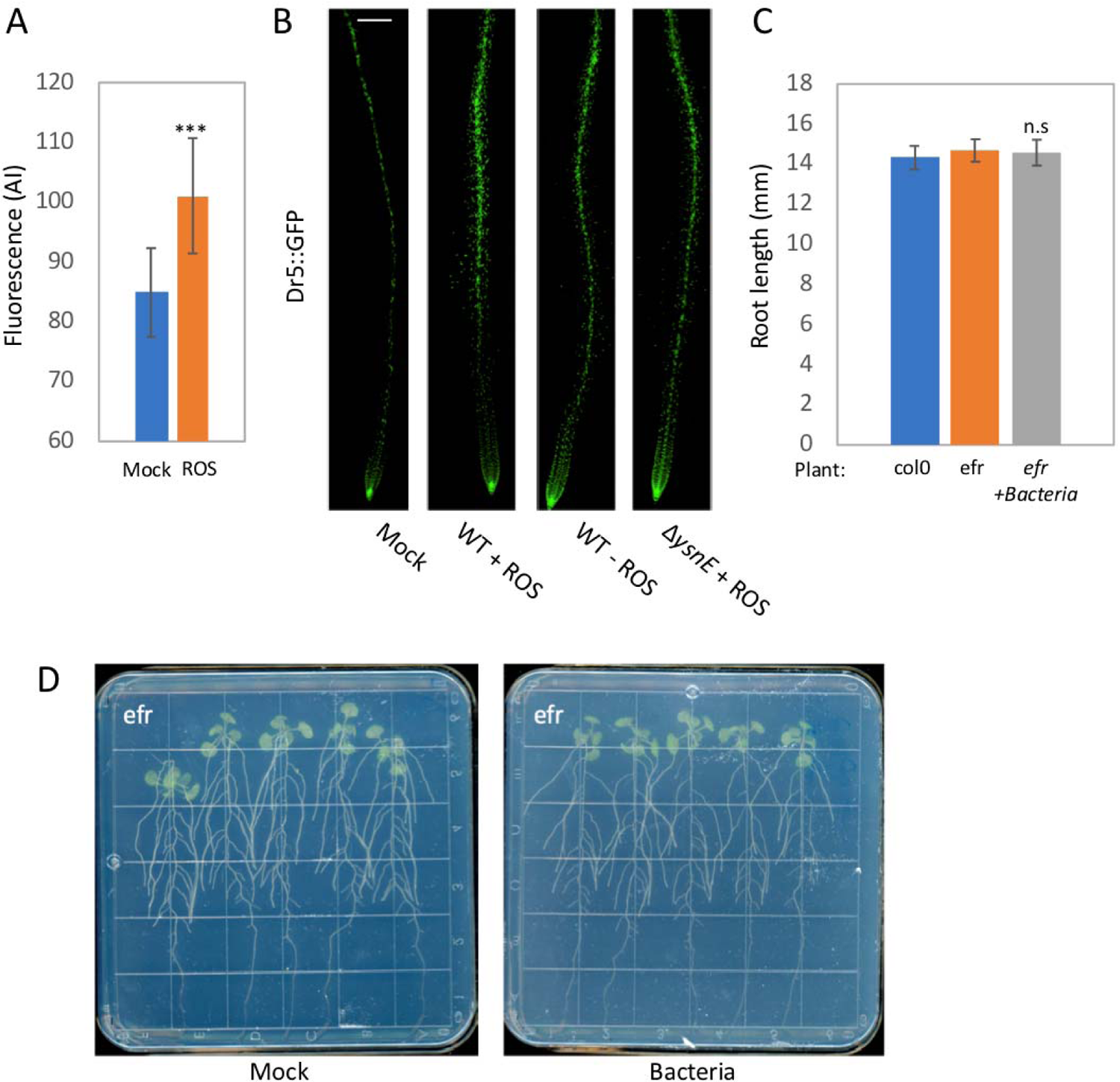
Plant immune system activation induces bacterial auxin secretion. (A) Ysne-GFP expressing bacteria grown to OD_600_=1 were treated with O^-^ for 30min and fluorescent intensity quantified. Shown are average bacterial fluorescence and SD of 4 independent fields, n ≥ 10 for each field. *** = *P <* 0.005. **(B)** Arabidopsis DR5::GFP reporter lines were inoculated for 12hrs with media derived from WT or *ΔysnE* bacteria grown to OD_600_=1 and treated with O^-^ through enzymatic reaction with xanthine and xanthine oxidase enzyme for 30min. Shown are 10x confocal images of DR5::GFP expression. Scale bar 50μm **(C)** Seedlings of col-0 or *efr2* were inoculated with bacteria or buffer alone (mock) on agar plates for 7 days and length of the primary root was measured, n ≥ 20. **(D)** *efr2* seedlings were inoculated with bacteria or buffer alone (mock) on agar plates for 7 days. Shown are plates from each of the conditions tested.

**Fig S6.**
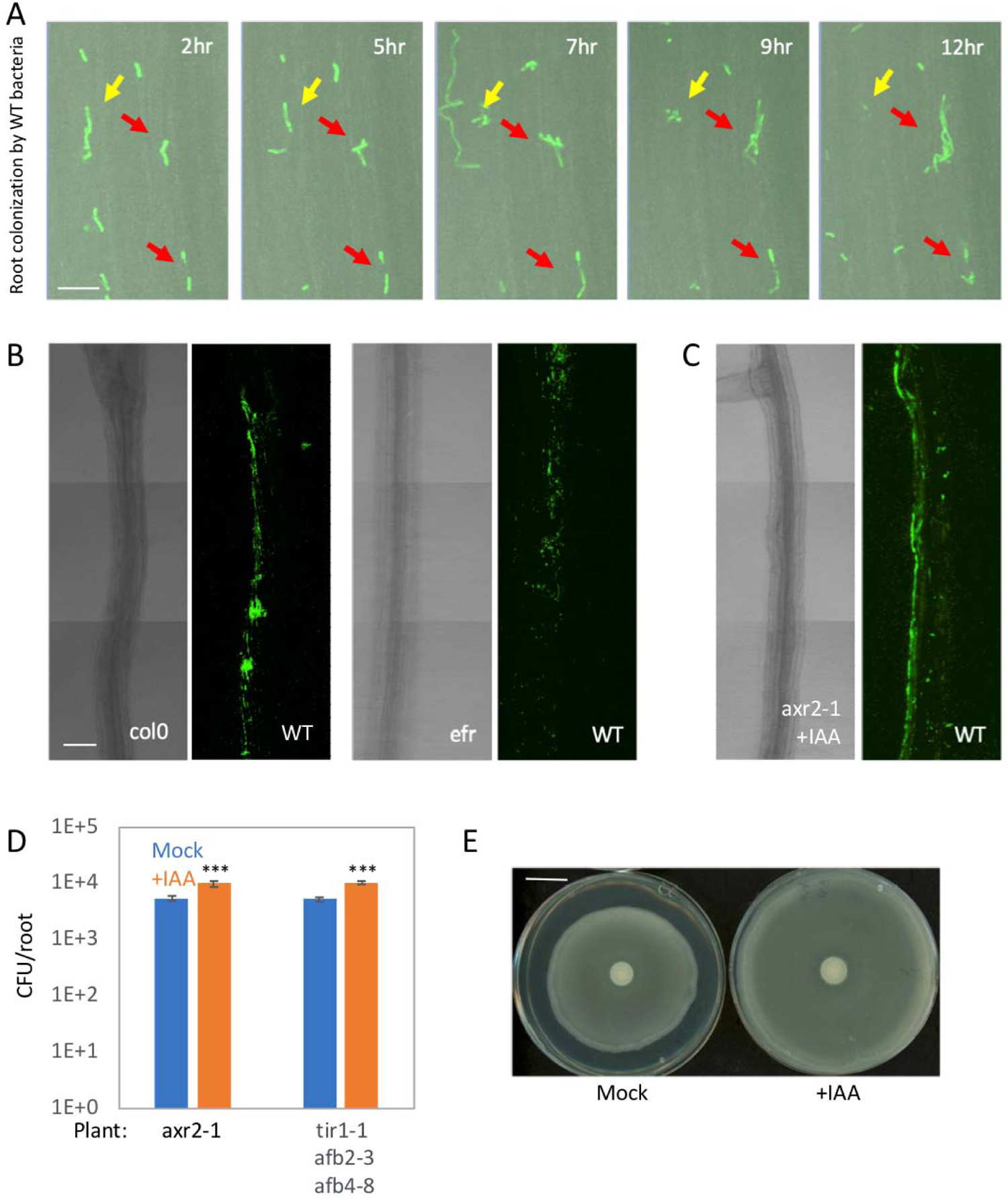
ROS induced auxin enhances root colonization through stimulation of bacterial flagella. (**A**) Col-0 or *efr2* seedling were inoculated with GFP expressing bacteria (*amyE::Pspac-gfp*) and then followed by time lapse confocal microscopy for 12hrs. Spots of colonizing bacteria were counted at t = 2hr and followed until t =12. Bacteria that remained attached during this time course were counted as successful colonization events (see Fig 4A for quantification). Shown are 400x overlay images of DIC from roots (grey) and GFP fluorescence from bacteria (green), taken at the indicated time points from *efr2* roots. Red arrows highlight bacteria counted as having undergone a successful colonization. Yellow arrows highlight bacteria counted as having undergona an unsuccessful colonization. Scale bar 10μm (**B-C**) Seedlings of the indicated genotypes were inoculted with WT bacteria expressing GFP, for 48 hrs on agar plates. In the absence (B) or presence of 5μM IAA (C). Shown are 200x confocal images of DIC from the roots (left panels) and GFP fluorescence from bacteria (right panels). Scale bar 50μm (D) Seedlings of the indicated genotypes were inoculated with WT bacteria in the presence or absence of 5μM IAA for 48 hrs on agar plates and the number of colonizing bacteria was counted. Shown are averages and SD from 2 independent experiments (log_10_ transformed) with n ≥ 3 for each. *** = *P <* 0.005. (E) Bacteria were inoculated in the middle of 0.7% agar plates with 5μM IAA or without (mock) and incubated at 37° for 9 hrs. Shown are representative plates from 3 plates for each condition. Scale bar 2mm.

**Fig S7.**
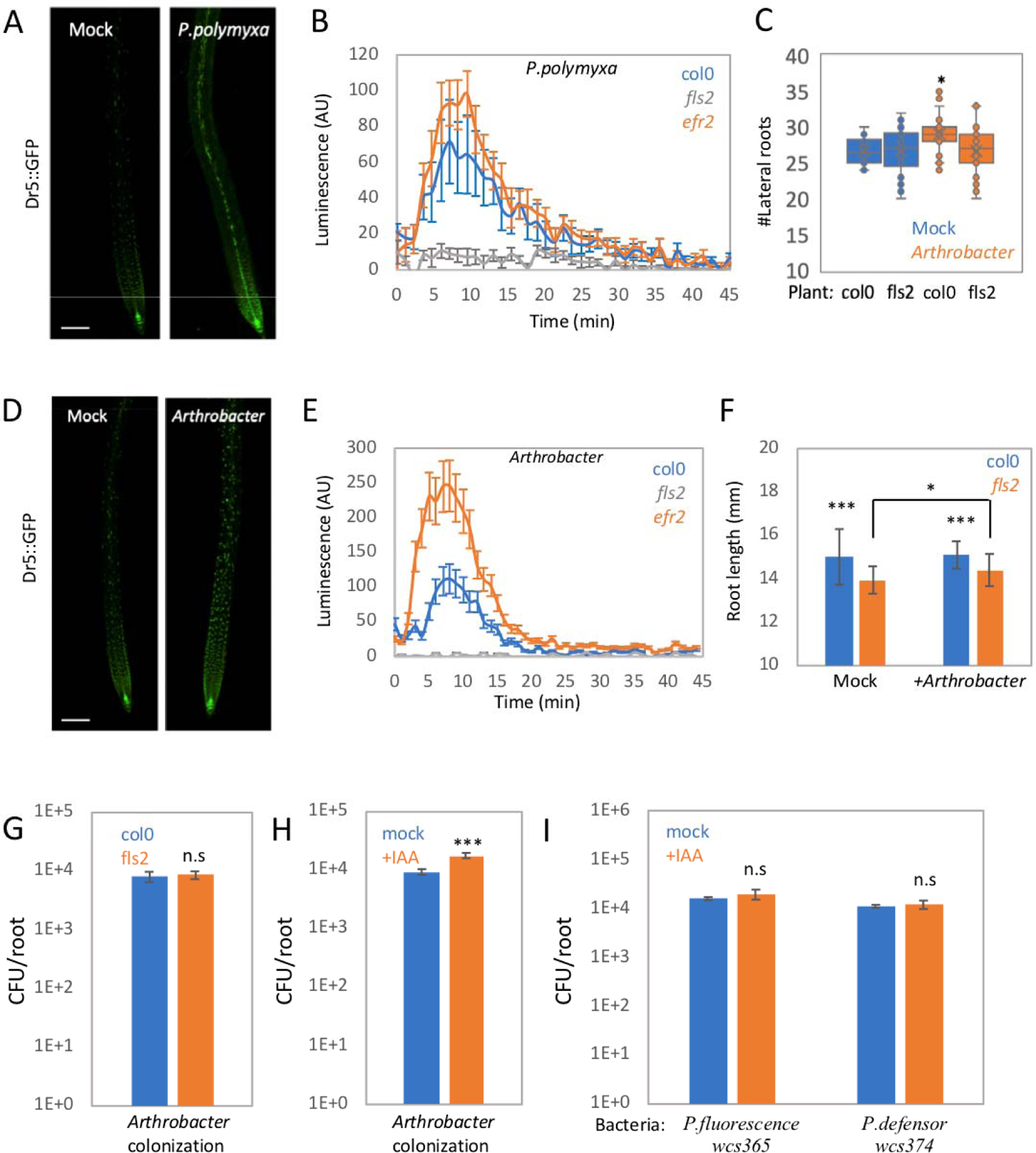
Plant immunity interaction with bacterial auxin secretion in *Arthrobacter mf161*. (A) Arabidopsis DR5::GFP reporter lines were inoculated with *P. polymyxa* for 96 hrs on agar plates. Shown are 100x confocal images of GFP fluorescence from DR5::GFP reporter line. (B) Leaf discs from 28 day-old plants, taken from the indicated plant genotypes were incubated with bacteria adjusted to OD 0.1, and ROS burst was measured. Shown are average and SD, n ≥ 10. (C) Col-0 or *fls2* seedlings were inoculated with *Arthrobacter mf161* or buffer alone (mock) on agar plates for 7 days and the number of lateral roots was counted, n 20. * = *P <* 0.05. (D) Arabidopsis DR5::GFP reporter lines were inoculated with *Arthrobacter mf161* or buffer alone (mock) for 48 hrs on agar plates. Shown are 100x confocal images of GFP fluorescence from DR5::GFP reporter. Scale bar 50μm. (E) Leaf discs from 28 day-old plants, taken from the indicated genotypes were incubated with bacteria adjusted to OD 0.1, and ROS production was measured. Shown are average and SD (n ≥ 10). (F) Col-0 or *fls2* seedlings were inoculated with *Arthrobacter mf161* or buffer alone (mock) on agar plates for 7 days and the length of the primary root was measured. (n 20) *** = *P <* 0.005. * = *P <* 0.05. (G) Col-0 or *fls2* seedlings were inoculated with *P. polymyxa* for 48 hrs on agar plates and the number of colonizing bacteria counted. Shown are averages and SD of 2 independent replicates (log_10_ transformed) with n = 3. (H) Seedlings were incubated with *P. polymyxa* for 48 hrs on agar plates in the presence or absence of 5μM IAA and the number of colonizing bacteria counted. Shown are averages and SD of 2 independent replicates (log_10_ transformed) with n=3 *** = *P < 0.005*. (I) Seedlings were incubated with *Pseudomonas fluorescence WCS365* or *Pseudomonas defensor WCS374* for 48 hrs on agar plates in the presence or absence of 5μM IAA and the number of colonizing bacteria counted. Shown are averages and SD (log_10_ transformed), n = 4. for each treatment.

**Fig S8.**
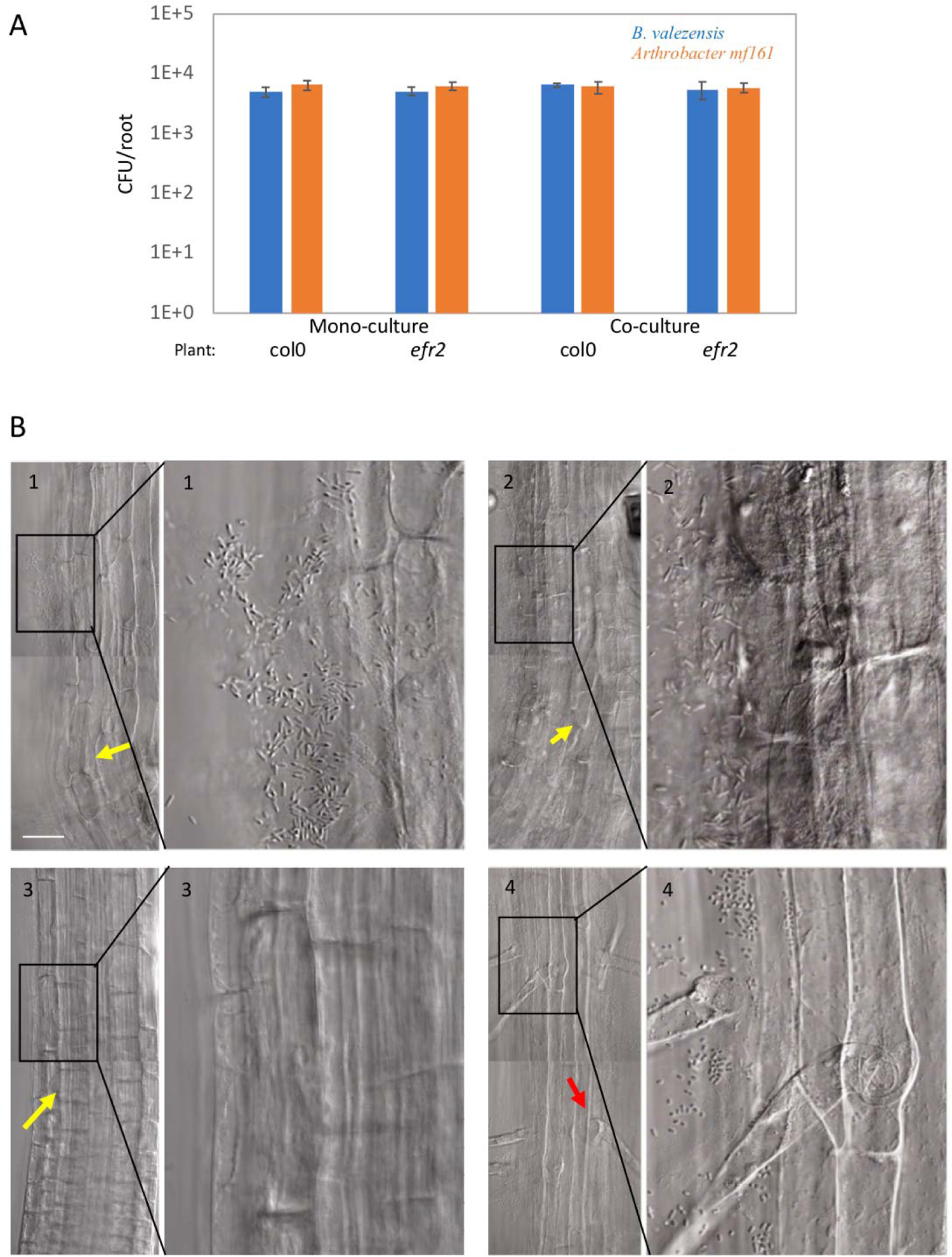
Immune system activation enhances bacterial colonization in face of competition. (A) Seedlings of Col-0 or *efr2* were inoculated with either *Arthrobacter mf161* or *B. valezensis* alone (monoculture) or in a mixture (1:1 ratio, co-culture) for 48 hrs on agar plates and the number of colonizing bacteria from each strain was counted. Shown are averages and SD of 2 independent replicates (log_10_ transformed) with n=3. (B) Seedlings were inoculated with the indicated bacterial strains for 48 hrs on agar plates. Shown are 400x confocal images of DIC from roots incubated with *B. valezensis* (1), *P. polymyxa* (2), and *Arthrobacter mf161* (3-4). The images on the left are magnifications of the black frames on the right images, in which the rod-shape *B. valezensis* (1) and *P. polymyxa* (2) and the small rod / cocci shape of the *Arthrobacter mf161* (4) can be seen. Yellow arrows highlight the small cells of the root elongation zone. Red arrow highlights the root hair of a trichoblast cell in the differentiated part of the root. Scale bar 10μm.

**Fig S9.**
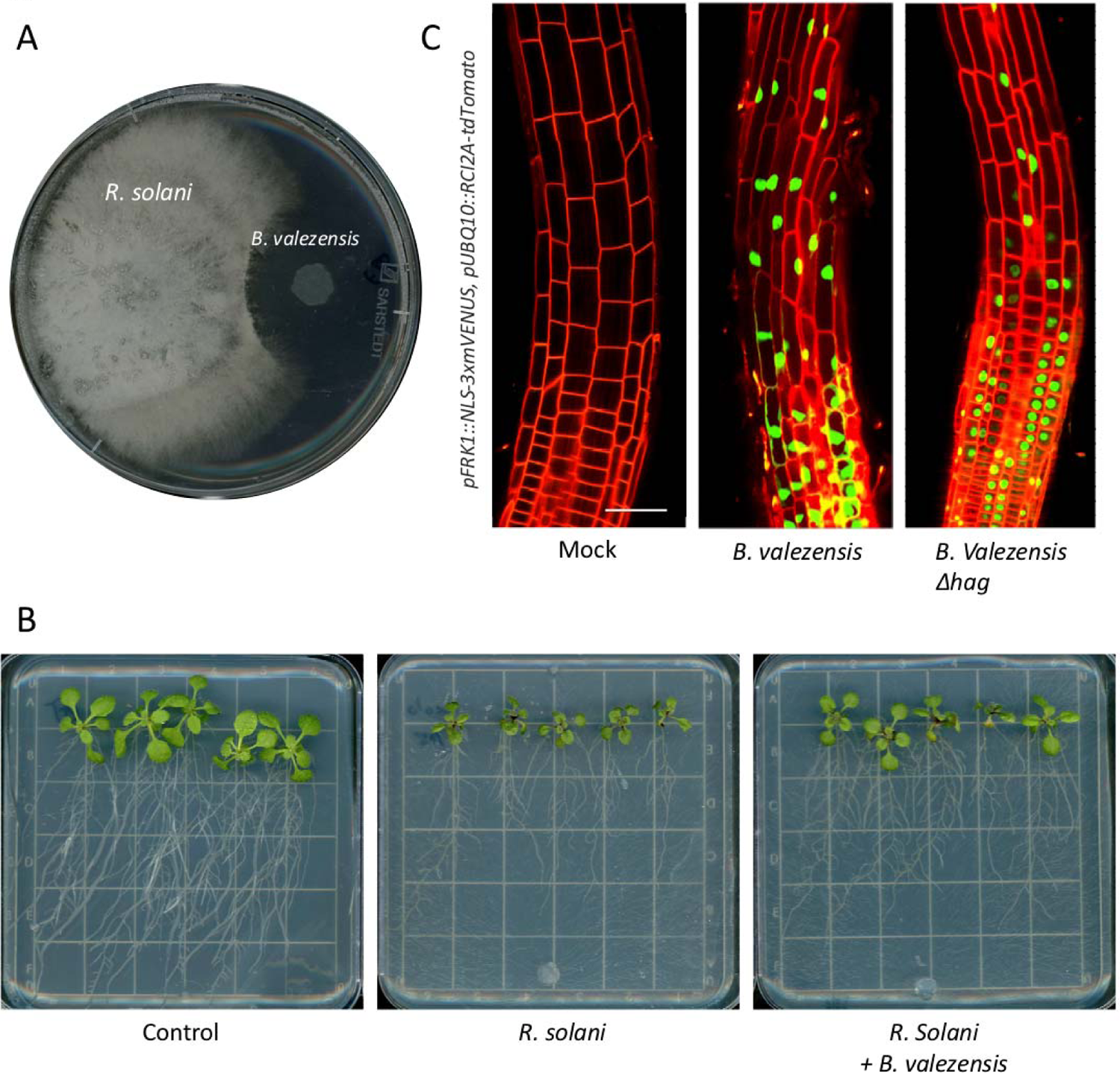
*B. valezensis* inhibits the growth of the fungal pathogen *R. solani*. **(A)** B. valezensis and R. solani were spotted on PDA plates and allowed to grow for 72 hrs. Shown is a representative plate from 3 plates. **(B)** 6 day-old seedlings were inoculated with *B. valezensis* or buffer for 48 hrs on agar plates. Then plates were inoculated with *R. solani* and incubated for an additional 7 days. Untreated plants were used as a control. Shown are representative plates from at least 5 plates for each treatment. **(C)** *pPER5::NLS-3xmVENUS*, *pUBQ10::RCI2A-tdTomato* seedlings were inoculated with either WT or *Δhag B. valezensis* or buffer alone (mock) for 48 hrs. Shown are 400x overlay images of *pUBQ10::RCI2A-tdTomato* (red) and *pPER5::NLS-3xmVENUS* (green) from 5 roots from each condition. Scale bars 25μm

## References

1. Lundberg, D. S. et al. Defining the core Arabidopsis thaliana root microbiome. Nature 488, 86–90, doi:10.1038/nature11237 (2012).

2. Bai, Y. et al. Functional overlap of the Arabidopsis leaf and root microbiota. Nature 528, 364–369, doi:10.1038/nature16192 (2015).

3. Couto, D. & Zipfel, C. Regulation of pattern recognition receptor signalling in plants. Nat Rev Immunol 16, 537–552, doi:10.1038/nri.2016.77 (2016).

4. Jones, J. D. & Dangl, J. L. The plant immune system. Nature 444, 323–329, doi:10.1038/nature05286 (2006).

5. Zipfel, C. Plant pattern-recognition receptors. Trends Immunol 35, 345–351, doi:10.1016/j.it.2014.05.004 (2014).

6. Zipfel, C. Early molecular events in PAMP-triggered immunity. Curr Opin Plant Biol 12, 414–420, doi:10.1016/j.pbi.2009.06.003 (2009).

7. Spoel, S. H. & Dong, X. How do plants achieve immunity? Defence without specialized immune cells. Nat Rev Immunol 12, 89–100, doi:10.1038/nri3141 (2012).

8. Dodds, P. N. & Rathjen, J. P. Plant immunity: towards an integrated view of plant-pathogen interactions. Nat Rev Genet 11, 539–548, doi:10.1038/nrg2812 (2010).

9. Xin, X. F., Kvitko, B. & He, S. Y. Pseudomonas syringae: what it takes to be a pathogen. Nat Rev Microbiol 16, 316–328, doi:10.1038/nrmicro.2018.17 (2018).

10. Stringlis, I. A. et al. Root transcriptional dynamics induced by beneficial rhizobacteria and microbial immune elicitors reveal signatures of adaptation to mutualists. Plant J 93, 166–180, doi:10.1111/tpj.13741 (2018).

11. Hacquard, S., Spaepen, S., Garrido-Oter, R. & Schulze-Lefert, P. Interplay Between Innate Immunity and the Plant Microbiota. Annu Rev Phytopathol 55, 565–589, doi:10.1146/annurev-phyto-080516-035623 (2017).

12. Teixeira, P. J. P., Colaianni, N. R., Fitzpatrick, C. R. & Dangl, J. L. Beyond pathogens: microbiota interactions with the plant immune system. Curr Opin Microbiol 49, 7–17, doi:10.1016/j.mib.2019.08.003 (2019).

13. Chen, T. et al. A plant genetic network for preventing dysbiosis in the phyllosphere. Nature 580, 653–657, doi:10.1038/s41586-020-2185-0 (2020).

14. Chowdhury, S. P. et al. Cyclic Lipopeptides of Bacillus amyloliquefaciens subsp. plantarum Colonizing the Lettuce Rhizosphere Enhance Plant Defense Responses Toward the Bottom Rot Pathogen Rhizoctonia solani. Mol Plant Microbe Interact 28, 984–995, doi:10.1094/MPMI-03-15-0066-R (2015).

15. Fan, B. et al. Bacillus velezensis FZB42 in 2018: The Gram-Positive Model Strain for Plant Growth Promotion and Biocontrol. Front Microbiol 9, 2491, doi:10.3389/fmicb.2018.02491 (2018).

16. Zhao, Y. Auxin biosynthesis and its role in plant development. Annu Rev Plant Biol 61, 49–64, doi:10.1146/annurev-arplant-042809-112308 (2010).

17. Banda, J. et al. Lateral Root Formation in Arabidopsis: A Well-Ordered LRexit. Trends Plant Sci 24, 826–839, doi:10.1016/j.tplants.2019.06.015 (2019).

18. Idris, E. E., Iglesias, D. J., Talon, M. & Borriss, R. Tryptophan-dependent production of indole-3-acetic acid (IAA) affects level of plant growth promotion by Bacillus amyloliquefaciens FZB42. Mol Plant Microbe Interact 20, 619–626, doi:10.1094/MPMI-20-6-0619 (2007).

19. Chowdhury, S. P. et al. Effects of Bacillus amyloliquefaciens FZB42 on lettuce growth and health under pathogen pressure and its impact on the rhizosphere bacterial community. PLoS One 8, e68818, doi:10.1371/journal.pone.0068818 (2013).

20. Fan, B. et al. Efficient colonization of plant roots by the plant growth promoting bacterium Bacillus amyloliquefaciens FZB42, engineered to express green fluorescent protein. J Biotechnol 151, 303–311, doi:10.1016/j.jbiotec.2010.12.022 (2011).

21. Mashiguchi, K. et al. Agrobacterium tumefaciens Enhances Biosynthesis of Two Distinct Auxins in the Formation of Crown Galls. Plant Cell Physiol 60, 29–37, doi:10.1093/pcp/pcy182 (2019).

22. Spaepen, S., Bossuyt, S., Engelen, K., Marchal, K. & Vanderleyden, J. Phenotypical and molecular responses of Arabidopsis thaliana roots as a result of inoculation with the auxin-producing bacterium Azospirillum brasilense. New Phytol 201, 850–861, doi:10.1111/nph.12590 (2014).

23. Kunkel, B. N. & Harper, C. P. The roles of auxin during interactions between bacterial plant pathogens and their hosts. J Exp Bot 69, 245–254, doi:10.1093/jxb/erx447 (2018).

24. Patten, C. L. & Glick, B. R. Bacterial biosynthesis of indole-3-acetic acid. Can J Microbiol 42, 207–220, doi:10.1139/m96-032 (1996).

25. Duca, D., Lorv, J., Patten, C. L., Rose, D. & Glick, B. R. Indole-3-acetic acid in plant-microbe interactions. Antonie Van Leeuwenhoek 106, 85–125, doi:10.1007/s10482-013-0095-y (2014).

26. Liao, C. Y. et al. Reporters for sensitive and quantitative measurement of auxin response. Nat Methods 12, 207–210, 202 p following 210, doi:10.1038/nmeth.3279 (2015).

27. Chen, Y. et al. Biocontrol of tomato wilt disease by Bacillus subtilis isolates from natural environments depends on conserved genes mediating biofilm formation. Environ Microbiol 15, 848–864, doi:10.1111/j.1462-2920.2012.02860.x (2013).

28. Dietel, K., Beator, B., Budiharjo, A., Fan, B. & Borriss, R. Bacterial Traits Involved in Colonization of Arabidopsis thaliana Roots by Bacillus amyloliquefaciens FZB42. Plant Pathol J 29, 59–66, doi:10.5423/PPJ.OA.10.2012.0155 (2013).

29. McClerklin, S. A. et al. Indole-3-acetaldehyde dehydrogenase-dependent auxin synthesis contributes to virulence of Pseudomonas syringae strain DC3000. PLoS Pathog 14, e1006811, doi:10.1371/journal.ppat.1006811 (2018).

30. Zhou, F. et al. Co-incidence of Damage and Microbial Patterns Controls Localized Immune Responses in Roots. Cell 180, 440–453 e418, doi:10.1016/j.cell.2020.01.013 (2020).

31. Zipfel, C. et al. Bacterial disease resistance in Arabidopsis through flagellin perception. Nature 428, 764–767, doi:10.1038/nature02485 (2004).

32. Zipfel, C. et al. Perception of the bacterial PAMP EF-Tu by the receptor EFR restricts Agrobacterium-mediated transformation. Cell 125, 749–760, doi:10.1016/j.cell.2006.03.037 (2006).

33. Willmann, R. et al. Arabidopsis lysin-motif proteins LYM1 LYM3 CERK1 mediate bacterial peptidoglycan sensing and immunity to bacterial infection. Proc Natl Acad Sci U S A 108, 19824–19829, doi:10.1073/pnas.1112862108 (2011).

34. Wu, Y. et al. Genome-Wide Expression Pattern Analyses of the Arabidopsis Leucine-Rich Repeat Receptor-Like Kinases. Mol Plant 9, 289–300, doi:10.1016/j.molp.2015.12.011 (2016).

35. Li, J. et al. Specific ER quality control components required for biogenesis of the plant innate immune receptor EFR. Proc Natl Acad Sci U S A 106, 15973–15978, doi:10.1073/pnas.0905532106 (2009).

36. Frerigmann, H., Berger, B. & Gigolashvili, T. bHLH05 is an interaction partner of MYB51 and a novel regulator of glucosinolate biosynthesis in Arabidopsis. Plant Physiol 166, 349–369, doi:10.1104/pp.114.240887 (2014).

37. Mucha, S. et al. The Formation of a Camalexin Biosynthetic Metabolon. Plant Cell 31, 2697–2710, doi:10.1105/tpc.19.00403 (2019).

38. Ding, P. & Ding, Y. Stories of Salicylic Acid: A Plant Defense Hormone. Trends Plant Sci 25, 549–565, doi:10.1016/j.tplants.2020.01.004 (2020).

39. Robert-Seilaniantz, A., Grant, M. & Jones, J. D. Hormone crosstalk in plant disease and defense: more than just jasmonate-salicylate antagonism. Annu Rev Phytopathol 49, 317–343, doi:10.1146/annurev-phyto-073009-114447 (2011).

40. Torres, M. A., Jones, J. D. & Dangl, J. L. Pathogen-induced, NADPH oxidase-derived reactive oxygen intermediates suppress spread of cell death in Arabidopsis thaliana. Nat Genet 37, 1130–1134, doi:10.1038/ng1639 (2005).

41. Tsukagoshi, H., Busch, W. & Benfey, P. N. Transcriptional regulation of ROS controls transition from proliferation to differentiation in the root. Cell 143, 606–616, doi:10.1016/j.cell.2010.10.020 (2010).

42. Fones, H. & Preston, G. M. Reactive oxygen and oxidative stress tolerance in plant pathogenic Pseudomonas. FEMS Microbiol Lett 327, 1–8, doi:10.1111/j.1574-6968.2011.02449.x (2012).

43. Wang, W. et al. Role of plant respiratory burst oxidase homologs in stress responses. Free Radic Res 52, 826–839, doi:10.1080/10715762.2018.1473572 (2018).

44. Engelmann, S. & Hecker, M. Impaired oxidative stress resistance of Bacillus subtilis sigB mutants and the role of katA and katE. FEMS Microbiol Lett 145, 63–69, doi:10.1111/j.1574-6968.1996.tb08557.x (1996).

45. Poole, L. B. Bacterial defenses against oxidants: mechanistic features of cysteine-based peroxidases and their flavoprotein reductases. Arch Biochem Biophys 433, 240–254, doi:10.1016/j.abb.2004.09.006 (2005).

46. Alonso, J. C. et al. Early steps of double-strand break repair in Bacillus subtilis. DNA Repair (Amst) 12, 162–176, doi:10.1016/j.dnarep.2012.12.005 (2013).

47. Cornelis, P., Wei, Q., Andrews, S. C. & Vinckx, T. Iron homeostasis and management of oxidative stress response in bacteria. Metallomics 3, 540–549, doi:10.1039/c1mt00022e (2011).

48. Chen, X. H. et al. Comparative analysis of the complete genome sequence of the plant growth-promoting bacterium Bacillus amyloliquefaciens FZB42. Nat Biotechnol 25, 1007–1014, doi:10.1038/nbt1325 (2007).

49. Jeong, H. et al. Draft genome sequence of the Paenibacillus polymyxa type strain (ATCC 842T), a plant growth-promoting bacterium. J Bacteriol 193, 5026–5027, doi:10.1128/JB.05447-11 (2011).

50. Castrillo, G. et al. Root microbiota drive direct integration of phosphate stress and immunity. Nature 543, 513–518, doi:10.1038/nature21417 (2017).

51. Kamilova, F., et al. Organic acids, sugars, and L-tryptophane in exudates of vegetables growing on stonewool and their effects on activities of rhizosphere bacteria. Mol Plant Microbe Interact 19, 250–256, doi:10.1094/MPMI-19-0250 (2006).

52. Zamioudis, C., Mastranesti, P., Dhonukshe, P., Blilou, I. & Pieterse, C. M. Unraveling root developmental programs initiated by beneficial Pseudomonas spp. bacteria. Plant Physiol 162, 304–318, doi:10.1104/pp.112.212597 (2013).

53. Dean, R. et al. The Top 10 fungal pathogens in molecular plant pathology. Mol Plant Pathol 13, 414–430, doi:10.1111/j.1364-3703.2011.00783.x (2012).

54. Pieterse, C. M. et al. Induced systemic resistance by beneficial microbes. Annu Rev Phytopathol 52, 347–375, doi:10.1146/annurev-phyto-082712-102340 (2014).

55. Cao, Y., Halane, M. K., Gassmann, W. & Stacey, G. The Role of Plant Innate Immunity in the Legume-Rhizobium Symbiosis. Annu Rev Plant Biol 68, 535–561, doi:10.1146/annurev-arplant-042916-041030 (2017).

56. Sachs, J. L., Skophammer, R. G. & Regus, J. U. Evolutionary transitions in bacterial symbiosis. Proc Natl Acad Sci U S A 108 Suppl 2, 10800–10807, doi:10.1073/pnas.1100304108 (2011).

57. Delaux, P. M. & Schornack, S. Plant evolution driven by interactions with symbiotic and pathogenic microbes. Science 371, doi:10.1126/science.aba6605 (2021).

58. Toth, K. & Stacey, G. Does plant immunity play a critical role during initiation of the legume-rhizobium symbiosis? Front Plant Sci 6, 401, doi:10.3389/fpls.2015.00401 (2015).

59. Miyata, K. et al. The bifunctional plant receptor, OsCERK1, regulates both chitin-triggered immunity and arbuscular mycorrhizal symbiosis in rice. Plant Cell Physiol 55, 1864-1872, doi:10.1093/pcp/pcu129 (2014).

60. Teale, W. D., Paponov, I. A. & Palme, K. Auxin in action: signalling, transport and the control of plant growth and development. Nat Rev Mol Cell Biol 7, 847–859, doi:10.1038/nrm2020 (2006).

61. Costacurta, A. & Vanderleyden, J. Synthesis of phytohormones by plant-associated bacteria. Crit Rev Microbiol 21, 1–18, doi:10.3109/10408419509113531 (1995).

62. Spaepen, S., Vanderleyden, J. & Remans, R. Indole-3-acetic acid in microbial and microorganism-plant signaling. FEMS Microbiol Rev 31, 425-448, doi:10.1111/j.1574-6976.2007.00072.x (2007).

63. Bianco, C. et al. Indole-3-acetic acid improves Escherichia coli’s defences to stress. Arch Microbiol 185, 373–382, doi:10.1007/s00203-006-0103-y (2006).

64. Van Puyvelde, S. et al. Transcriptome analysis of the rhizosphere bacterium Azospirillum brasilense reveals an extensive auxin response. Microb Ecol 61, 723–728, doi:10.1007/s00248-011-9819-6 (2011).

65. Djami-Tchatchou, A. T. et al. Dual Role of Auxin in Regulating Plant Defense and Bacterial Virulence Gene Expression During Pseudomonas syringae PtoDC3000 Pathogenesis. Mol Plant Microbe Interact 33, 1059–1071, doi:10.1094/MPMI-02-20-0047-R (2020).

66. Zhang, P. et al. The Distribution of Tryptophan-Dependent Indole-3-Acetic Acid Synthesis Pathways in Bacteria Unraveled by Large-Scale Genomic Analysis. Molecules 24, doi:10.3390/molecules24071411 (2019).

67. Finkel, O. M. et al. A single bacterial genus maintains root growth in a complex microbiome. Nature 587, 103–108, doi:10.1038/s41586-020-2778-7 (2020).

68. Guan, G. et al. PfeT, a P1B4-type ATPase, effluxes ferrous iron and protects Bacillus subtilis against iron intoxication. Mol Microbiol 98, 787–803, doi:10.1111/mmi.13158 (2015).

69. Rosenberg, A., Sinai, L., Smith, Y. & Ben-Yehuda, S. Dynamic expression of the translational machinery during Bacillus subtilis life cycle at a single cell level. PLoS One 7, e41921, doi:10.1371/journal.pone.0041921 (2012).

70. Branda, S. S., Gonzalez-Pastor, J. E., Ben-Yehuda, S., Losick, R. & Kolter, R. Fruiting body formation by Bacillus subtilis. Proc Natl Acad Sci U S A 98, 11621–11626, doi:10.1073/pnas.191384198 (2001).

71. Schikora, S. T. S. A. Staining of Callose Depositions in Root and Leaf Tissues bio-protocol, doi:https://doi.org/10.21769/BioProtoc.1429 (2015).

72. Bray, N. L., Pimentel, H., Melsted, P. & Pachter, L. Near-optimal probabilistic RNA-seq quantification. Nat Biotechnol 34, 525–527, doi:10.1038/nbt.3519 (2016).

73. Wachsman, G., Zhang, J., Moreno-Risueno, M. A., Anderson, C. T. & Benfey, P. N. Cell wall remodeling and vesicle trafficking mediate the root clock in Arabidopsis. Science 370, 819–823, doi:10.1126/science.abb7250 (2020).

